# Spatial integration of mechanical forces by alpha-actinin establishes actin network symmetry

**DOI:** 10.1101/578799

**Authors:** Fabrice Senger, Amandine Pitaval, Hajer Ennomani, Laetitia Kurzawa, Laurent Blanchoin, Manuel Théry

## Abstract

Cell and tissue morphogenesis depend on the production and spatial organization of tensional forces in the actin cytoskeleton. Actin network architecture is complex because it is made of distinct modules in which filaments adopt a variety of organizations. The assembly and dynamics of these modules is well described but the self-organisation rules directing the global network architecture are much less understood. Here we investigated the mechanism regulating the interplay between network architecture and the geometry of cell’s extracellular environment. We found that α-actinin, a filament crosslinker, is essential for network symmetry to be consistent with extracellular microenvironment symmetry. It appeared to be required for the interconnection of transverse arcs with radial fibres to ensure an appropriate balance between forces at cell adhesions and across the entire actin network. Furthermore, the connectivity of the actin network appeared necessary for the cell ability to integrate and adapt to complex patterns of extracellular cues as they migrate. Altogether, our study has unveiled a role of actin-filament crosslinking in the physical integration of mechanical forces throughout the entire cell, and the role of this integration in the establishment and adaptation of intracellular symmetry axes in accordance with the geometry of extracellular cues.

## Introduction

Actin filaments self-organize into a variety of modules, and the combination of modules can generate a diversity of sub-cellular architectures (Blanchoin et al., 2014). These modules include lamellipodia, radial fibres, transverse arcs and stress fibres. The lamellipodium consists of branched and entangled filaments. Radial fibres consist of parallel bundles of filaments, which are anchored on focal adhesions and orientated radially toward cell centre. Transverse arcs and stress fibres are contractile modules consisting of anti-parallel bundles of filaments. Transverse arcs are localized in the lamella and align parallel to adhesive cell edges. Stress fibres connect two distant focal adhesions, and traverse the non-adhesive part of the cell periphery or the central part of the cell (Tojkander et al., 2012).

Actin-based modules are dynamic. In the lamellipodium and radial fibres, actin dynamic is dominated by actin polymerization, whereas in transverse arcs or stress fibres dynamic is governed by acto-myosin contraction. All undergo regulated disassembly. Growth, contraction and disassembly coexist in the cell, leading to complex intracellular patterns of expansion and shrinkage (Levayer and Lecuit, 2012; Belmonte et al., 2017). Furthermore, modules interconvert (Letort et al., 2015). Lamellipodium filaments can align to form transverse arcs when they are pushed away from their site of nucleation at the cell membrane (Burnette et al., 2011; Shemesh et al., 2009). Radial fibres and transverse arcs tend to join and coalesce to form stress fibres (Hotulainen and Lappalainen, 2006; Tojkander et al., 2015).

The combination of internal dynamics and interconversion of modules leads to a complex dynamic network architectures. Theoretically, this could lead to disordered phases (Mak et al., 2016). However, in cells, it results into surprisingly ordered and reproducible architectures. Yet, how modules consisting of self-organized filaments of 100 nm to 1 *µ*m in length can result in an architecture that is 10-to 100-fold larger with a geometry and symmetry that is in correspondence with the cell’s environment remains an open question.

The relative dispositions of regions containing growing and contracting modules define cell polarity. Indeed, intracellular signalling, and feedback between different signalling pathways, enforces defined spatial patterns of actin organization, through processes like mutual exclusion. This process of mutual exclusion prevents the assembly of a patchwork of modules throughout the cell, and primes the segregation of regions of growing modules and contracting modules and stabilizes their relative positioning (Lim et al., 2013; Chau et al., 2012). In addition, the determination of module types and their relative positioning is also finely regulated. Actin network self-organization accurately responds to extracellular cues. Chemical stimulation, extracellular adhesion, and the rigidity and geometry of the extracellular environment can regulate the self-organization of actin modules, thus conferring cells with the ability to sense and adapt to altered patterns of chemical or physical signals from the cellular environment (Young et al., 2016).

The multitude of different signals implies a process of signal integration, which results in a single polarity axis being defined and ensures that intracellular polarity is adapted to extracellular cues. For example, epithelial cells assemble specific actin-based architecture at their apical pole and thus orientate their polarity toward the lumen. Migrating cells assemble a characteristic network at the cell front guiding them along signal gradients. This integration process ensures the consistency between intracellular architecture and the geometry of the extracellular environment.

This spatial integration is the outcome of the self-organization of both signalling pathways (Chau et al., 2012) and cytoskeletal networks (Ingber, 2003; Fletcher and Mullins, 2010; Lee and Kumar, 2016; Vignaud et al., 2012a). Although the crosstalk between signalling pathways leading to cell polarization has been extensively studied (Lim et al., 2008), less attention has focussed on the integration of physical forces leading to ordered and regular intracellular architectures.

In terms of cytoskeletal networks, the spatial integration of extracellular cues is based on actin network connectivity throughout the cell. Localized damage to a cell leads to rapid relaxation/reorientation of actin-mediated forces far from the site of damage, in a time scale that is so brief that it precludes a process of signalling-pathway reconfiguration (Kumar et al., 2006; Chang and Kumar, 2013); and a single site of detachment of a cell from the extracellular matrix can affect the direction of cell motility (Verkhovsky et al., 1999). Hence these examples illustrate the existence of mechanical force transmission throughout the cell. Altogether these considerations suggest that the production of mechanical forces and their transmission throughout the actin network contribute to probe the extra-cellular environment and organize cell architecture accordingly. Such a working model assumes that the cell adhesion machinery translate the extracellular cues into a pattern of nucleating and anchoring sites for actin filaments; a pattern which is further reorganized by contractile forces. The transmission of forces throughout the entire actin network effectively acts as a process of integration allowing forces on all adhesion sites to be balanced within a single interconnected architecture. In addition, the contribution of contractile forces to network rearrangement, such as filament displacements and module inter-conversion, allows the final cell architecture to be geometrically and mechanically consistent with cell microenvironment. Notably, such a remodelling of the actin network by the transmission and balance of contractile forces applied on anchorage sites should allow this network to adopt the same elements of symmetry as the extracellular environment. Here, we interrogate this working model by investigating the architecture and symmetry of contractile actin networks in response to controlled geometrical cues.

## Results

### Crosslinkers support actin networks contraction symmetry in vitro

The architecture and polarity of the cellular actin network are shaped by the combined inputs from signalling pathways and structural self-organisation. However, the contributions of these respective inputs are difficult if not impossible to distinguish. The reconstitution of actin-network growth and contraction in vitro using purified components is an effective way to investigate network structural self-organization in a system that is independent of the spatial patterning of signalling pathways. Several recent examples illustrate the ability of reconstituted network to recapitulate key cellular actin network properties such as anchoring-dependent reinforcement (Ciobanasu et al., 2014), long range coherence (Ennomani et al., 2016b; Linsmeier et al., 2016), contraction-disassembly coupling (Murrell and Gardel, 2012; Reymann et al., 2012; Vogel et al., 2013), symmetry breaking (Abu Shah and Keren, 2014) and scaling of contractile rates (Reymann et al., 2012).

Recently, the spatial distribution of motors was shown to direct the shape changes of contracting actin networks (Schuppler et al., 2016). Interestingly, the asymmetric positioning of anchorage sites generated anisotropic network contraction (Schuppler et al., 2016). However, network deformation is the outcome of the mechanical work of motors on a given network architecture (Reymann et al., 2012). The role of actin-network architecture on the direction and symmetry of its contraction remains to be investigated. Here, we addressed this point by looking at the contraction of actin rings in vitro.

A given actin network architecture, such as a ring-shaped structure (actin ring), can be created by the surface micropatterning of actin nucleation sites (Reymann et al., 2010), and its contraction can be monitored following the addition of myosin (Ennomani et al., 2016b; Reymann et al., 2012). Surprisingly, with an actin ring, the pattern of contraction was heterogeneous and asymmetric. The initial bias in ring deformation was amplified leading to eccentric contraction. The focus point for the contraction was distant from the centre and nucleation pattern of the initial ring-shaped structure, and therefore appeared not to arise from contraction towards this centre (figure 1B, D, movie S1)

**Figure 1:**
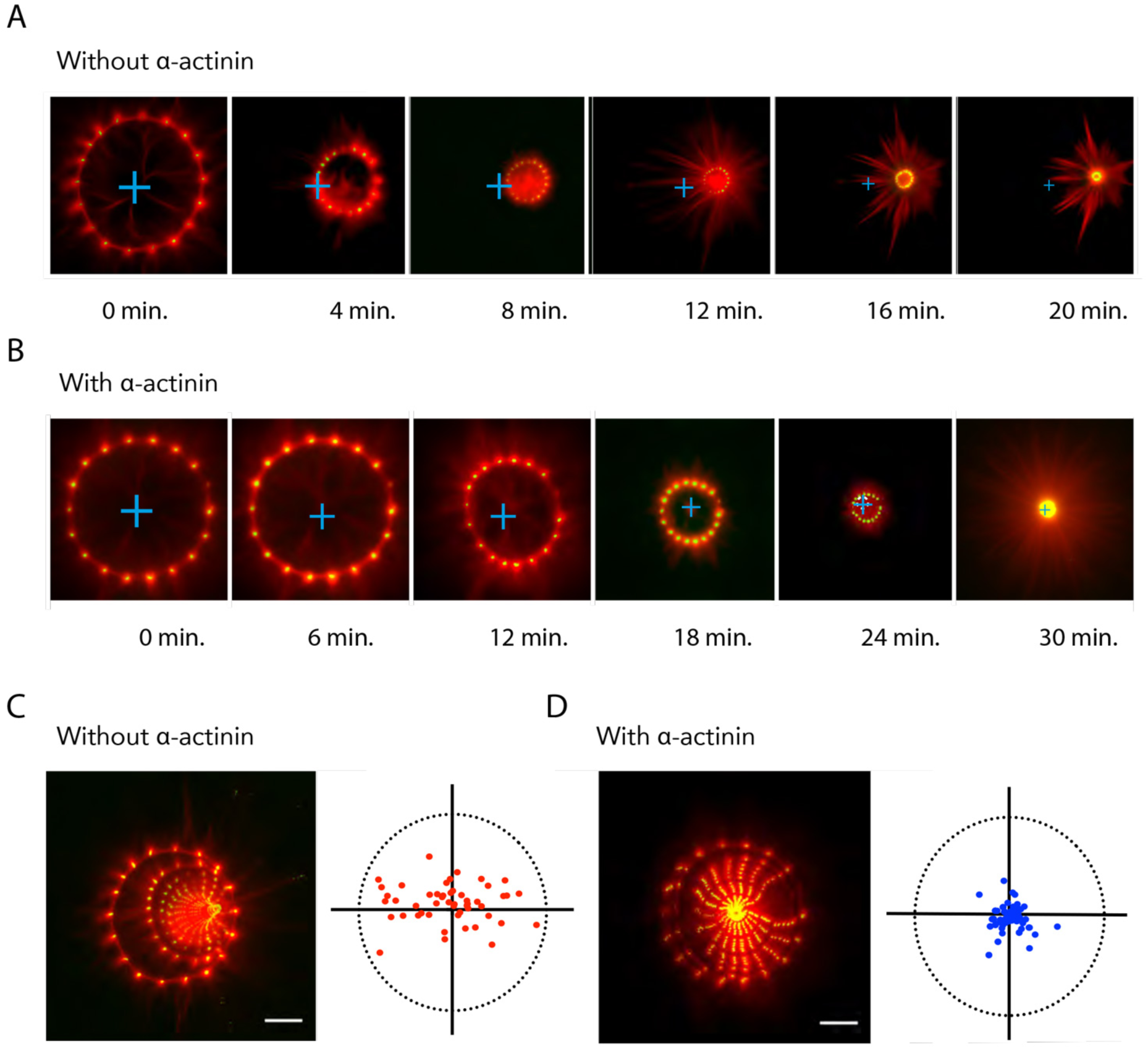
Alpha-actinin modulates the symmetry of actin network contractility *in-vitro.* Actin-ring assembly stemmed from the polymerization of actin filament from micropatterned dots coated with actin nucleation promoting factors. (A, B) contraction of the actin rings upon addition of myosin motors in the absence (A) or presence (B) of α-actinin. Time is indicated in minutes. t=0 corresponds to the onset of contraction. (C, D) Images show the maximal projection of pictures showed in (A) and (B) respectively. The plots represent the coordinates of the centre of the contractile ring after full contraction (C, n=73; D, n=50). Scale bar represents 20 *µ*m.

Crosslinkers are key regulator of actin network contractility (Jensen et al., 2015; Ennomani et al., 2016b) and are likely to be required for local contraction to be transmitted throughout the network by controlling the network connectivity. Strikingly, the addition of the filament crosslinker, α-actinin, fully restored network contraction symmetry. Actin rings displayed more progressive and regular contraction. The initial ring-shaped structure appeared to be conserved throughout the contractile process. And the final focus point of the contraction was close to the centre of the nucleation pattern (Figure 1A, C, movie S1).

### α-actinin knock-down impairs cellular actin network symmetry

These results prompted us to investigate whether actin network crosslinking was required for the transmission of mechanical forces throughout the cell and thereby for the proper integration of extracellular cues during the self-organization of cellular actin network architecture. Surface micropatterning was used to investigate cell responses to defined adhesion patterns with controlled elements of symmetry and anisotropy (Théry, 2010).

On crossbow-shaped patterns, cells developed a stereotypical actin architecture, presenting a branched network forming the lamellipodium along the curved adhesive edge, which further reorganized in the lamella to form radial fibres perpendicular to the curved edge and transverse arcs along the curved edge, as well as an array of ventral stress fibres above non-adhesive edges flanking the central bar of the crossbow pattern (Théry et al., 2006; Schiller et al., 2013; Jiu et al., 2015) (Figure 2A, movie S2).

**Figure 2:**
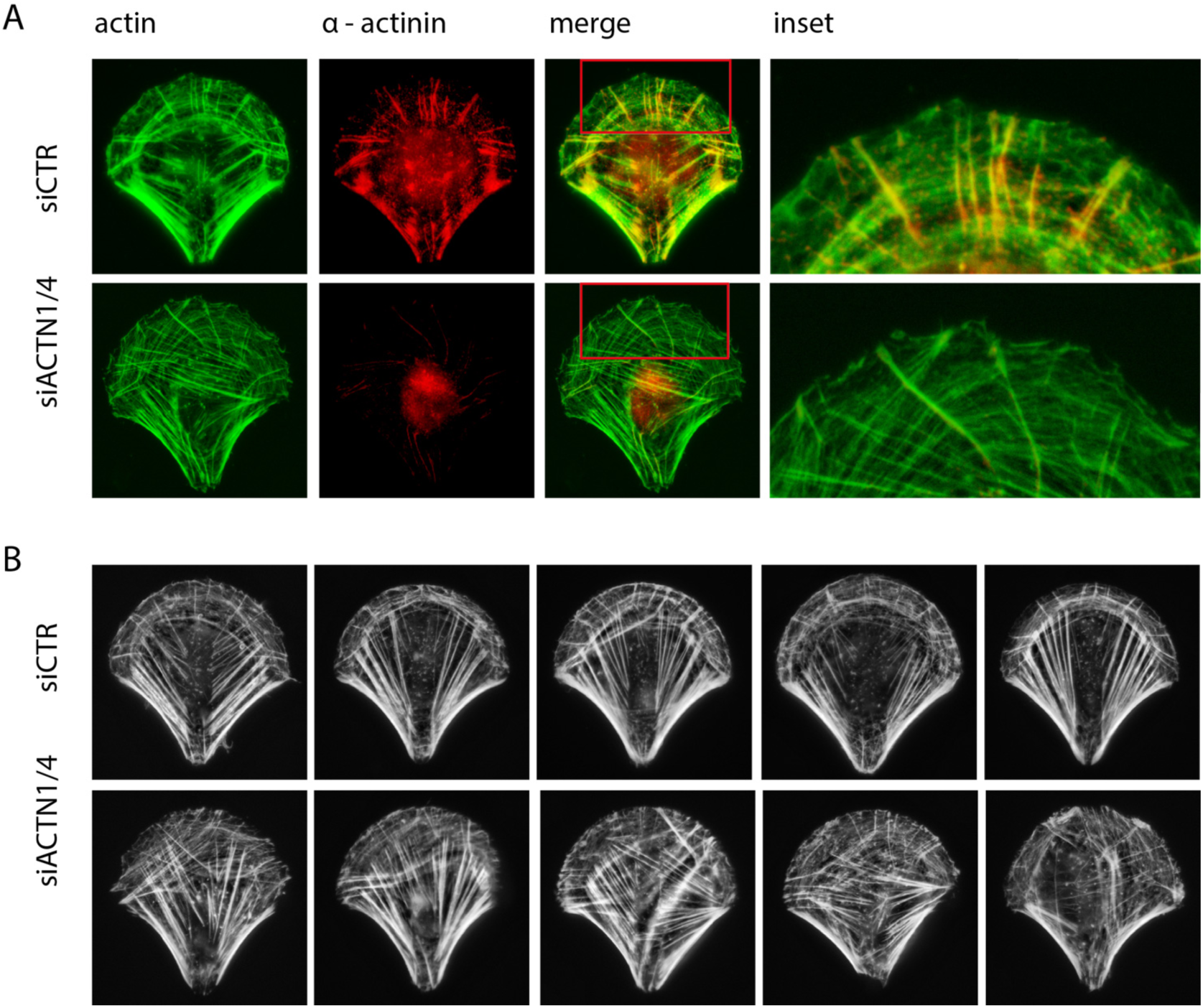
Reorganization of cellular actin architecture upon a-actinin depletion. (A) RPE1 cells on crossbow shaped micropatterns were fixed and stained for actin (green) and α-actinin (red). Inserts show a magnified view of the upper part of the cell and the loss of α-actinin staining upon α-actinin depletion (siACTN1/4, bottom row). (B) Phalloidin staining revealing F-actin architecture in control (siCTR) and α-actinin depleted cells (siACTN1/4).

Alpha-actinin localized at focal adhesions, along radial fibres and transverse arcs, and more particularly at the junction between the latter two structures (Figure 2A). Such localization at the intersections of the main contractile actin-based modules suggests a potential role in ensuring the connectivity of the entire network and the transmission of contractile forces across the cell body.

We attempted to identify the contribution of α-actinin to actin network architecture by down-regulating its expression with short interfering RNAs (siRNA). Alpha-actinin is present in two isoforms, alpha-actinin-1 (ACTN1) and alpha-actinin-4 (ACTN4). In RPE1 cells, ACTN1 expression level seems to compensate for ACTN4 down-regulation (Figure S1). Therefore a double knockdown against both isoforms appeared necessary to efficiently deplete the presence of α-actinins on the network (Figure S1). The specificity of our siRNA approach has been assessed by a redundancy check, i.e., the use of a second set of siRNA targeting another sequence of respectively ACTN1 and ACTN4 (Figure S2).

As a result of the double a-actinin knockdown, many cells had troubles to fully spread on the micropatterns. In spread cells the actin network appeared strikingly disorganised. Transverse arcs were diminished, if not absent, and radial fibres seemed misoriented (Figure 2B), as previously described (Kovac et al., 2013). Notably, the symmetrical organization of all types of actin bundles appeared to be compromised with respect to the symmetry axis of the fibronectin micropattern (Figure 3A).

**Figure 3:**
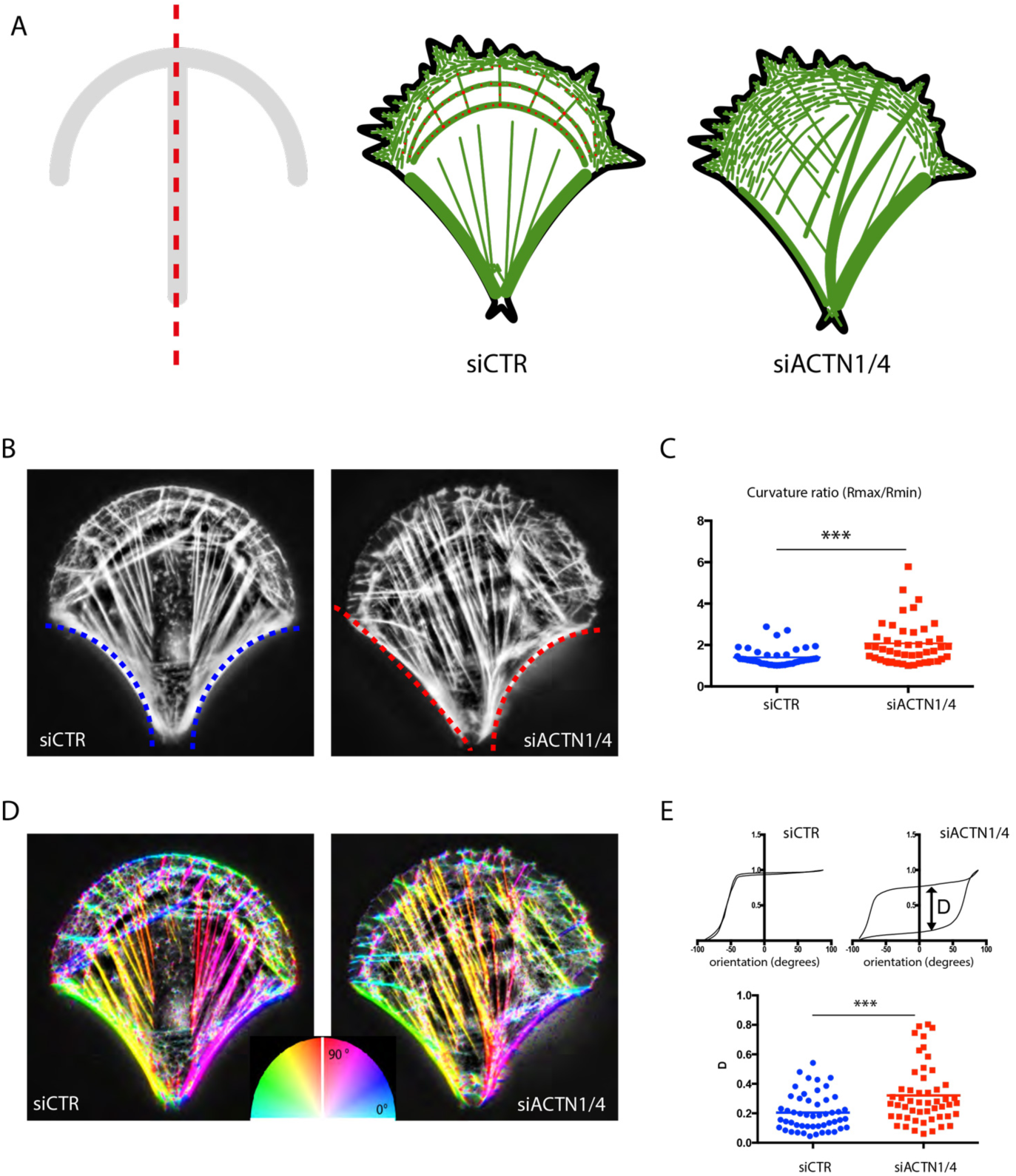
Loss of actin-network symmetry upon a-actinin depletion. (A) Scheme showing the symmetry axis of the micropattern coated with extracellular matrix (red dashed line) and the actin-network architecture in control (siCTR) and a-actinin depleted cells (siACTN1/4). (B) Measurement of the curvature of the two non-adherent edges in control (siCTR, blue fit) and a-actinin depleted cells (siACTN1/4, red fit). (C) The maximal curvature (Rmax) to minimal curvature (Rmin) ratio was calculated and plotted for the control (n=43) and a-actinin depleted cells (n=45). Statistical significance has been assessed by Mann-Whitney test (p < 0.0001). (D) Detection and measurement of internal actin-bundle orientation. Actin-fibre orientation is colour coded according to color-wheel inset. (E) Upper graphs show examples of cumulative distribution of actin-bundle angles as determined by orientation analysis for control and a-actinin depleted cells. In each graph, the two curves correspond to angular distributions of the left and the right half of the cell with respect to the micropattern symmetry axis. The two curves were compared by measuring their maximal difference D. Lower plots show the values of parameter D for control (n=51) and a-actinin depleted cells (n=52).

To quantify the structural consequence of a-actinin depletion on the actin network, we first compared the curvature radii of the two peripheral stress fibres. The curvature radii of the two stress fibres on both sides of the micropattern symmetry axis were almost identical in control conditions (with a median ratio of 1.274) and significantly different in α-actinin depleted cells (with a median ratio of 1.797) (Figure 3B, C). We then segmented and measured the orientation of actin bundles (Figure 3D). We compared the cumulative angular distributions away from the axis of micropattern symmetry of individual actin bundles in the two respective halves of the cell on each side of this axis (Figure 3E). In control conditions, the difference in these cumulative angular distributions between the two halves of the cell appeared much lower in the control conditions than after a-actinin depletion, and therefore appeared more in line with the axis of mirror symmetry (Figure 3E). To assess the specificity of alpha-actinin knock-down in these experiments, we performed rescue experiments which showed that cells recovered their symmetry upon expression of another alpha-actinin construct which was not sensitive to the set of siRNAs (Figure S3).

### Alpha-actinin knock-down deviates actin flow and nucleus position

By considering that actin bundles result from the conversion of lamellipodium fibres into transverse arcs and further into stress fibres as they move toward cell centre, it is likely that the misorientated actin bundles in α-actinin depleted cells were associated with a misorientated inward flow of contractile transverse arcs. Time-lapse recording of Lifeact-GFP expressing cells showed that actin bundles assembled near the curved adhesive edge were progressively displaced toward the cell centre over time (Figure 4A, movie S3). Image segmentation followed by particle-image velocimetry was then used to track the moving structures. In control cells, contractile actin bundles converged towards the cell centre, and hence the averaged flow vector was aligned with the micropattern symmetry axis. By contrast, in α-actinin depleted cells, the average flow vector deviated from the symmetry axis (Figure 4B, C). This observation was consistent with a recent observation of a bias in the actin flow at the immunological synapse in response to a-actinin depletion (Ashdown et al., 2017)

**Figure 4:**
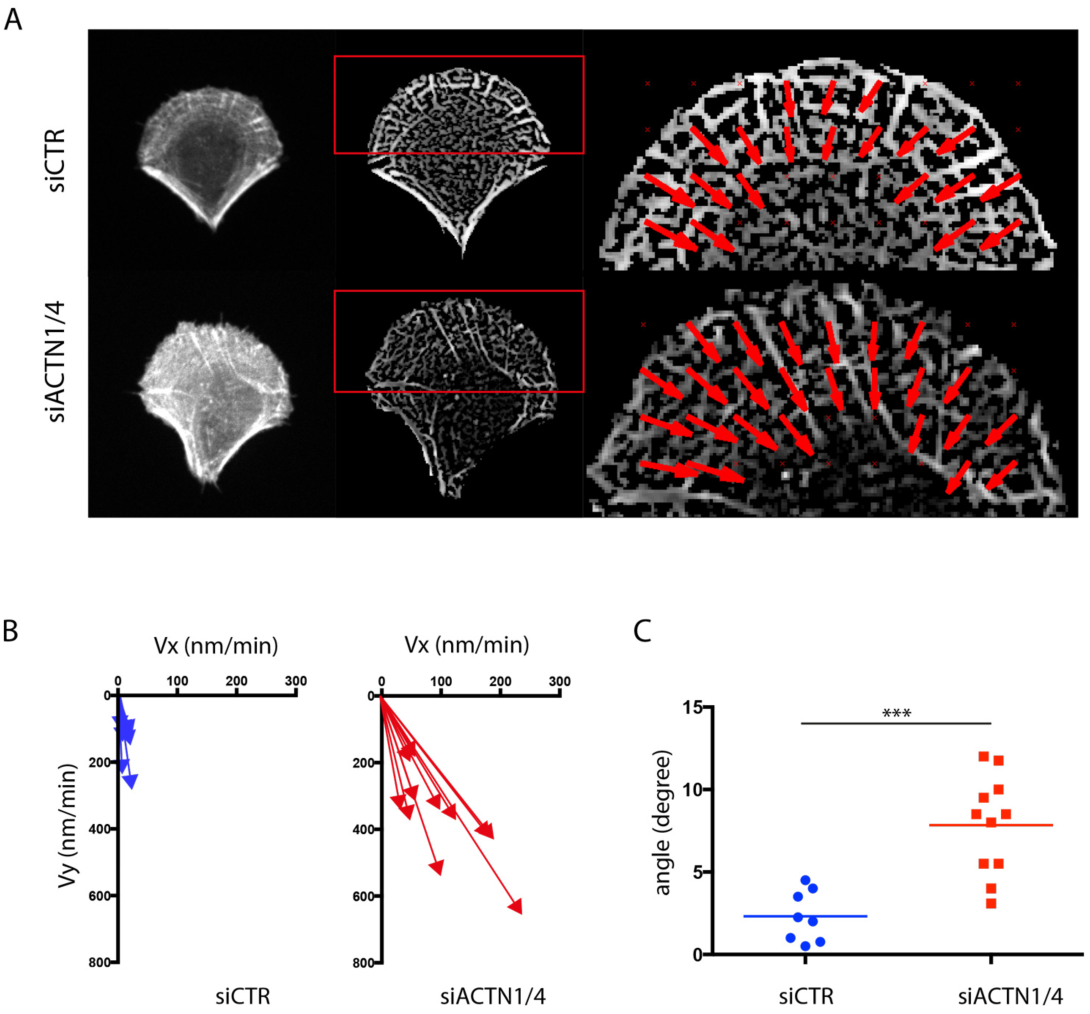
Misorientated actin network flow upon a-actinin depletion. (A) Images show LifeAct-GFP signal in live RPE1 cells plated on crossbow-shaped micropatterns (left), filtered images with noise removed (centre) and a zoomed image of the PIV flow on the upper part of the cell (right). Red arrows show the flow orientation. Statistical significance has been assessed by Mann-Whitney test (p = 0.0003). (B) Graphs show the orientations of the averaged flow in different cells. Arrows lengths correspond to the averaged flow speed and Vx and Vy to their cartesian coordinates. (C) Plots show the value of the angular deviation of the average flow with respect to the micropattern symmetry axis for control (n=8) and a-actinin depleted cells (n=11). Statistical significance has been assessed by Mann-Whitney test (p =
0.0003).

The retrograde flow of contractile-actin bundles is involved in nucleus positioning (Gomes et al., 2005) with which they interact. Asymmetric actin flow could therefore affect the proper positioning of the nucleus. In a control cell, the nucleus was positioned on the symmetry axis, near the cell centre (Figure 5A, B) whereas in an a-actinin depleted cell, the nucleus was positioned away from the cell centre towards one of the two peripheral stress fibres (Figure 5A, B). Comparison of the distributions of nucleus distance to the symmetry axis confirmed a significant deviation in actinin-depleted cells (Figure 5C). Hence these result supported the conclusion that a-actinin-mediated regulation of the retrograde flow of contractile-actin bundles played role in aligning the organization and the symmetry axis of the cell with that of the extracellular environment.

**Figure 5:**
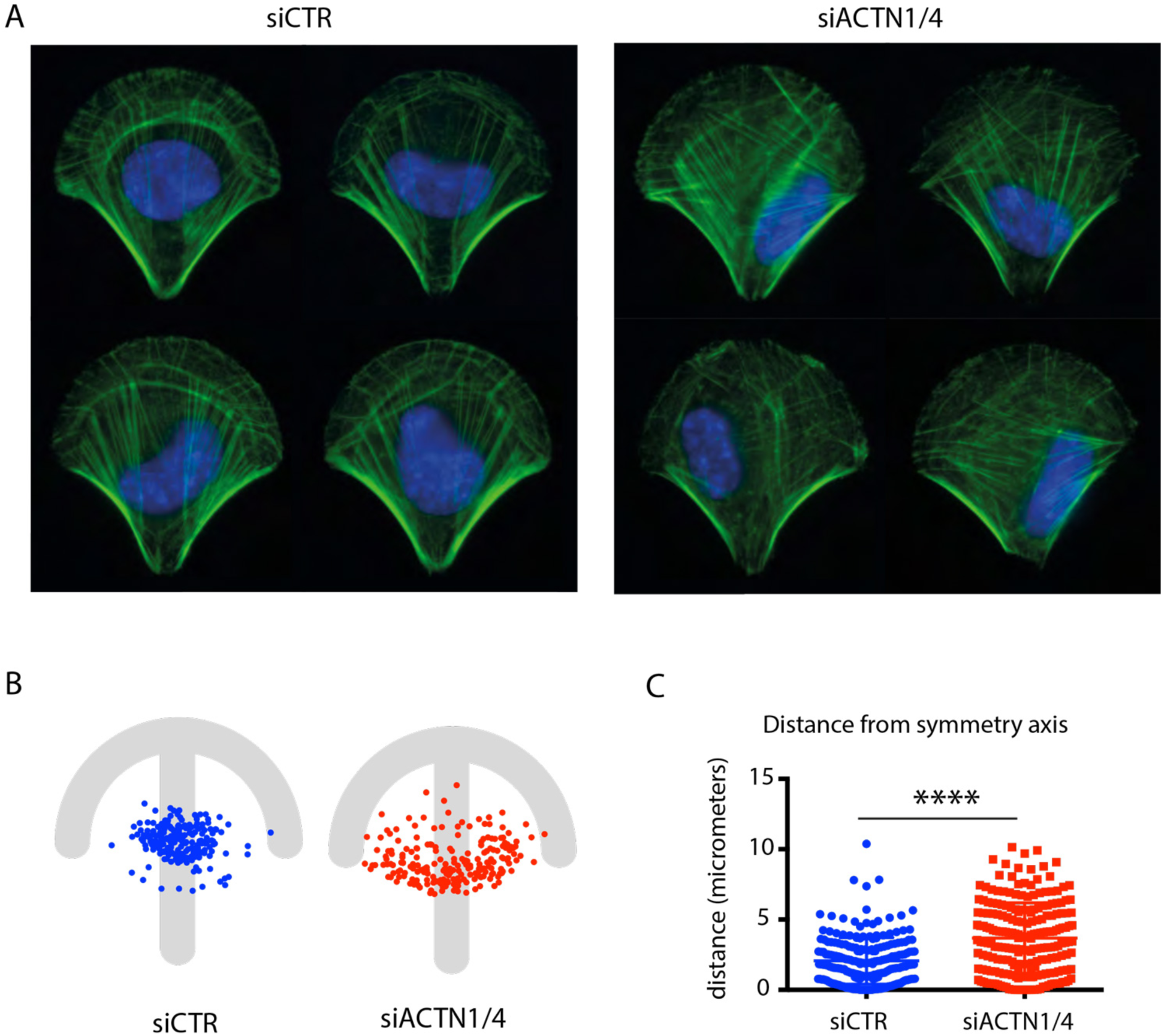
Nucleus mispositioning upon a-actinin depletion. (A) F-actin (green) and nuclei (DAPI, blue) stainings in control (siCTR) and a-actinin (siACTN1/4) depleted cells. (B) Spatial distribution of positions of centres of mass of nuclei in control (siCTR, blue, n=210) and a-actinin depleted cells (siACTN1/4, red, n=214) with respect to micropattern. (C) The plot shows the distance between the center of mass of the nucleus and the symmetry axis of the micropattern in control (siCTR, blue, n=210) and a-actinin depleted cells (siACTN1/4, red, n=214) (Mann_Whitney, p<0.0001).

### Asymmetric distribution of mechanical forces in α-actinin depleted cells

We then characterized the molecular machinery and the spatial distribution of mechanical forces associated with cell contractile modules. With respect to the molecular machinery, we examined the localization of non-muscle myosin IIA, IIB and the phosphorylated myosin light chains. As expected (Naumanen et al., 2008) these myosin components were found on transverse arcs and stress fibres. (Figure 6A). In accordance with the alignment of the actin modules with the axis of symmetry of the crossbow micropattern, the distributions of myosin components appeared precisely symmetrical with respect to this axis in control cells but significantly biased in α-actinin depleted cells (Figure 6B). The myosin-component asymmetry stemmed from the asymmetry of the two peripheral stress fibres as well as from defective transverse arcs. These transverse arcs were less bundled and misorientated compared with those in control cells in which they formed conspicuous and regular circular arcs lining the curved cell edge.

**Figure 6:**
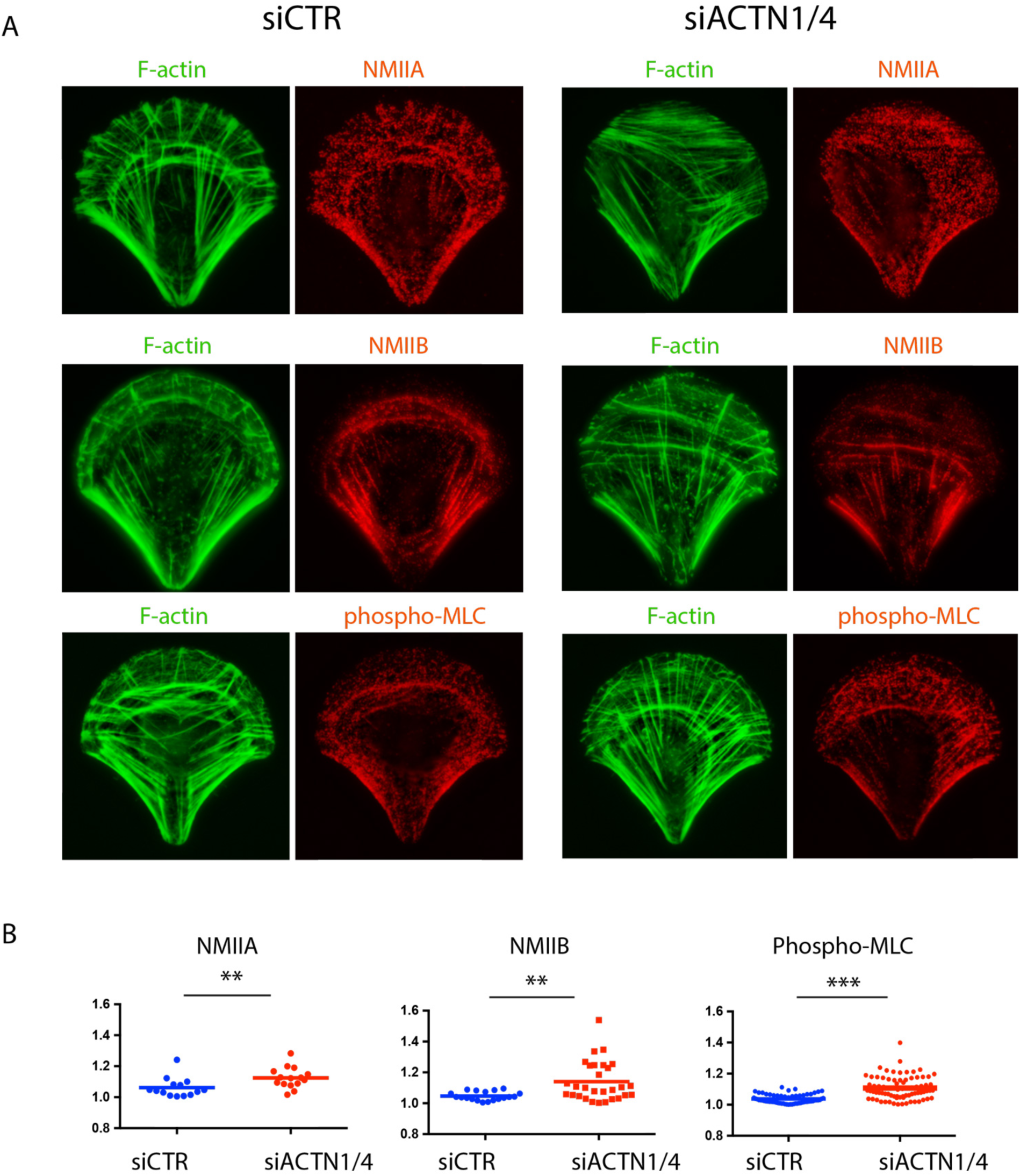
Myosin mislocalization upon a-actinin depletion. (A) F-actin (green) and myosins (red) stainings in control and a-actinin depleted cells. First row shows non-muscle myosin IIA (NMIIA), second row non-muscle myosin IIB (NMIIB) and third row phosphorylated myosin light chains (phospho-MLC). (B) Myosin fluorescence signal intensities were measured in each half of the cell with respect to the micropattern symmetry axis. Graphs show the ratio of the maximal to the minimal intensity. Non-muscle myosin IIA (NMIIA) (siCTR, n=14; siACTN1/4 = 15). Non-muscle myosin IIB (NMIIB) (siCTR, n=19; siACTN1/4 = 31). Phospho myosin light chain (P-MLC) (siCTR, n=63; siACTN1/4 = 79). Statistical significance has been assessed by Mann-Whitney test for NMIIA (p=0.0043), NMIIB (p = 0.0017) and P-MLC (p < 0.0001).

The contractile forces produced by these myosins also appeared asymmetrically distributed in α-actinin depleted cells. To measure these forces we used fibronectin micropatterns on polyacrylamide hydrogels in which fiduciary beads were embedded (Vignaud et al., 2014). Cells were first plated onto those micropatterns and then detached. The bead displacement field associated with hydrogel relaxation following cell detachment was monitored and computed to infer the associated traction force field (Martiel et al., 2015).

As expected (Tseng et al., 2011; Sun et al., 2016; Schiller et al., 2013), forces were evenly distributed along the curved adhesive edge, and were similarly concentrated at each extremity of the edge, and at the extremity of the central bar corresponding to the apex of the fan shaped distribution of contractile bundles (Figure 7A). In α-actinin depleted cells, the forces were less evenly distributed along the curved adhesive edge, and tended to be concentrated at only one of the two extremities of the edge (Figure 7A). The lower concentration of forces at the extremity of the edge was consistent with the defective assembly of transverse arcs and the misorientation of stress fibres. The main orientation of the traction force field was then characterized by measuring the first order moment of the force force field (Butler et al., 2002; Mandal et al., 2014).

**Figure 7:**
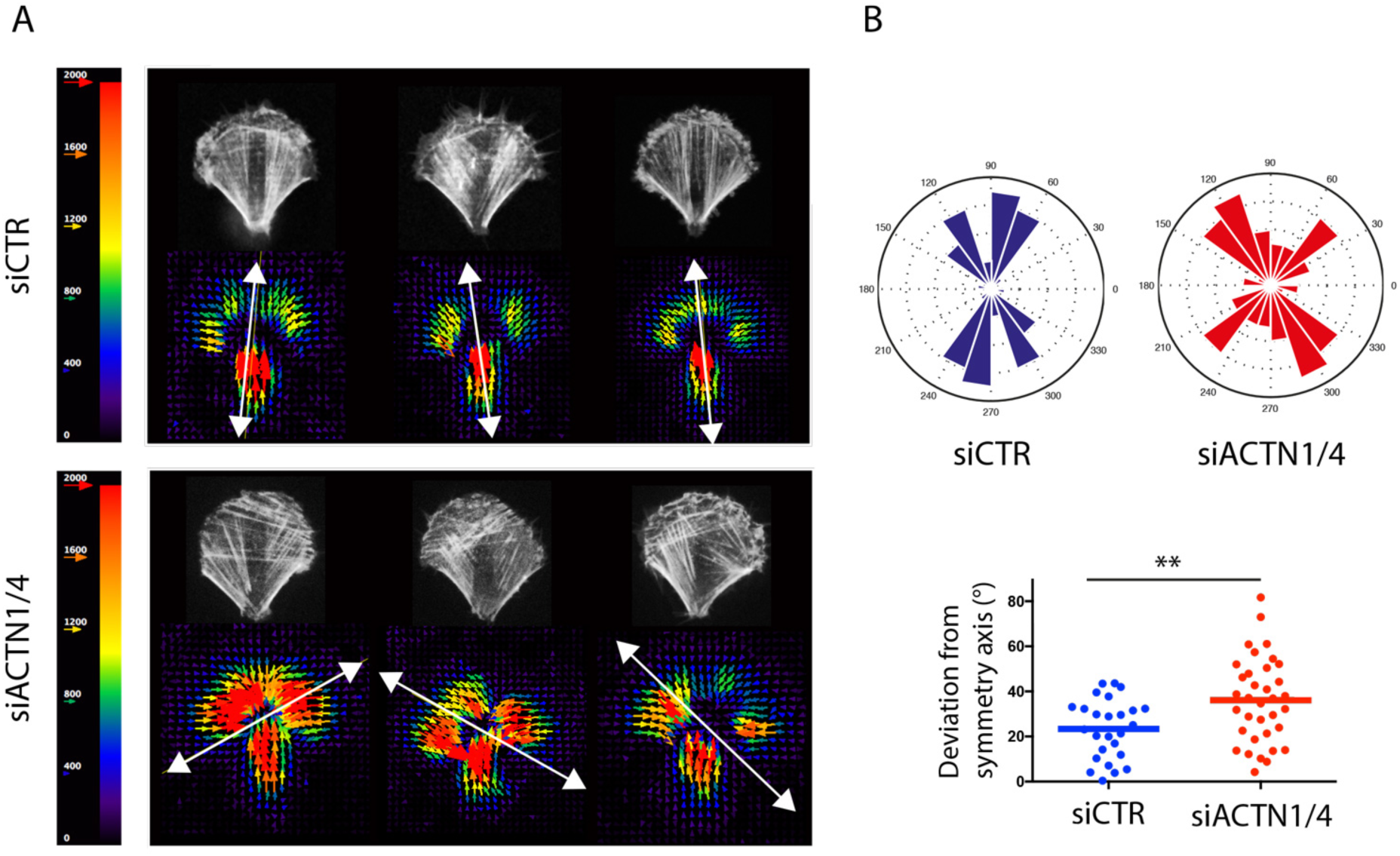
Traction force redistribution upon a-actinin depletion. (A) Images show traction force fields in control (siCTR; left) and a-actinin depleted cells (siACTN1/4; right). Arrows indicate force orientation and colour and length represent local stress magnitude in pascals (Pa). Force dipole calculation revealed the main axis within force field (white double headed arrow). (B) Rose-plot depicts the frequency distribution of the calculated force dipoles orientation for siCTR in blue (n=26); and siACTN1/4 in red (n=35). Graph depicts deviation (in degrees) from symmetry axis. Statistical significance has been assessed by Mann-Whithney test (p = 0.0097).

In control cells, the traction force field was generally aligned with the micropattern symmetry axis (Figure 7B). By contrast, in α-actinin depleted cells, it often appeared deflected from the symmetry axis reflecting the greater concentration of traction forces at only one of the two extremities of the curved adhesive edge (Figure 7B). Nevertheless, α-actinin depleted cells were able to produce traction forces, which were even stronger than the forces produced in control cells, as previously reported (Roca-Cusachs et al., 2013; Oakes et al., 2012). Some of the forces were aligned with the symmetry axis and appeared to be produced above non-adhesive regions by central or peripheral stress fibers. Therefore the forces driving the asymmetry in the traction force field may have been due to the forces produced in the lamella by the misorientated transverse arcs that were positioned at the curved adhesive edge and across the symmetry axis.

### Defective force transmission across the cell in α-actinin depleted cells

To characterize force transmission along the curved adhesive edge and perpendicular to the symmetry axis we looked more closely at the network architecture by analyzing the actin-network skeleton and by measuring forces relaxation in response to nano-ablation of subcellular structures.

The integrity of network architecture in the lamella relies on the connection of radial fibres with transverse arcs and on the redistribution of contractile forces produced in the arcs toward adhesion sites via radial fibres. The structural integrity of this module can be captured by measuring the overall connectivity of the actin network. Image segmentation of phalloidin-stained cells was used to skeletonize the actin network architecture (Figure 8A). The same threshold and segmentation parameters were applied to control and α-actinin depleted cells. The number of junctions and the length of connected bundles in the actin-network skeletons were significantly lower in α-actinin depleted cells than control cells confirming the lack of structural connectivity in these cells (Figure 8B).

**Figure 8:**
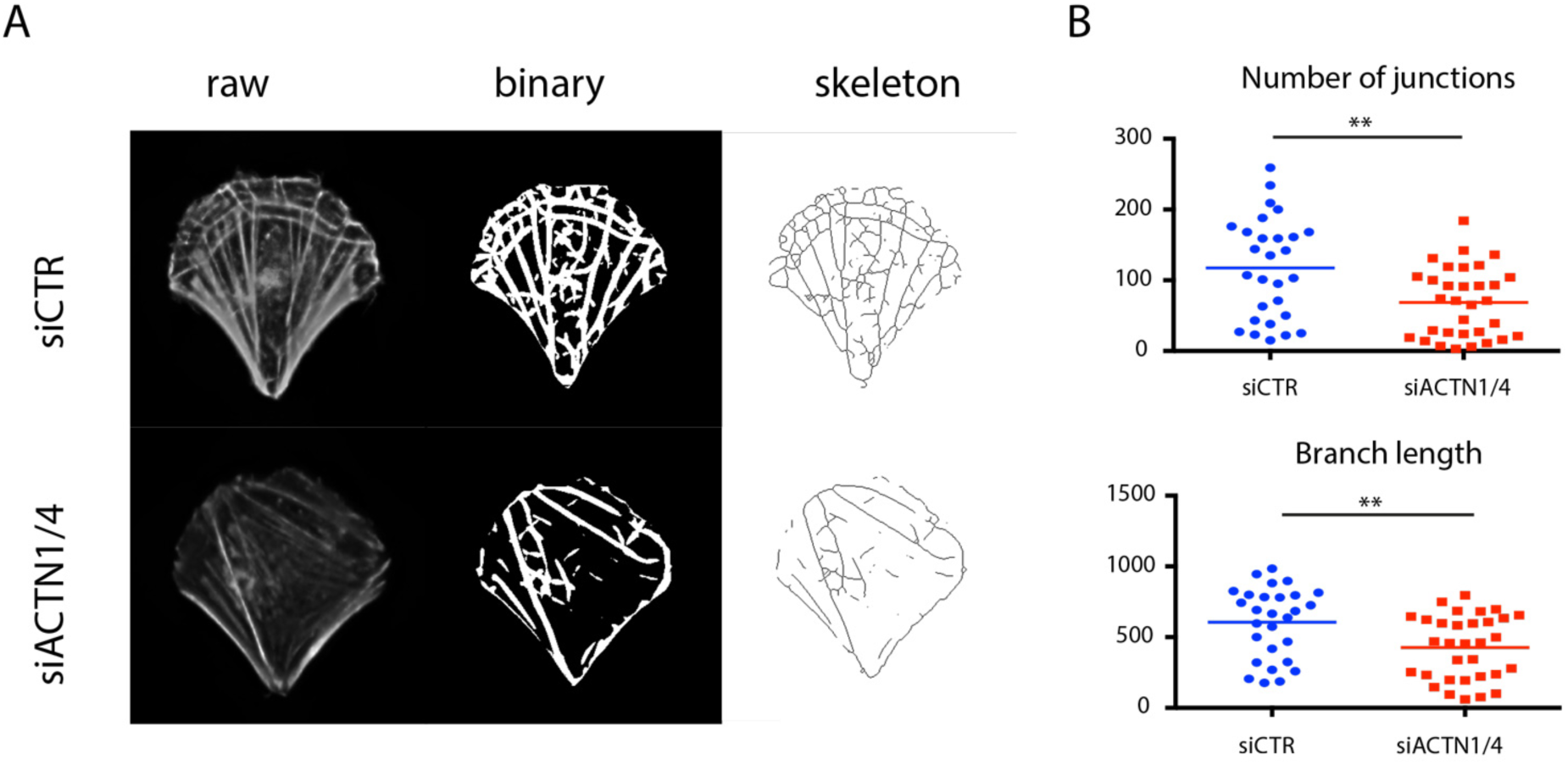
Lack of actin-network connectivity upon a-actinin depletion. (A) Images show F-actin staining in RPE1 cells plated on crossbow-shaped micropatterns (left), filtered, segmented and binarized images (centre) and the corresponding skeletons (right) in control (siCTR; top row) and a-actinin depleted cells (siACTN1/4; bottom row). (B) The number of junctions between branches were measured on skeletonized images and compared between control and a-actinin depleted cells. Length of skeleton corresponds to the product of average branch length times number of branches. (siCTR, n=28; siACTN1/4, n=32) (Statistical significance has been assessed by Mann-Whithney test. For junctions p=0.0042 and for Skeleton length, p = 0.0044).

The contribution to the global traction force field of tensional forces from transverse arcs and radial fibers was assessed by nano-ablation of subcellular structures positioned on the micropattern symmetry axis, 5 microns away from the curved adhesive edge (Figure 9A). The magnitude and orientation traction force field in ablated structures was inferred from hydrogel relaxation on which the cells were plated (Figure 9B, movie S4). In control cells, the ablated tensional forces produced by transverse arcs were transmitted perpendicular to the micropattern symmetry axis. By contrast, in α-actinin depleted cells, the ablated tensional forces aligned with the symmetry axis (Figure 9B). These results confirmed that the crosslinking by α-actinin of the actin network was necessary for the transmission of tensional forces across the lamella in the cell. In absence of α-actinin the various elements of the adhesive front of the cell were not properly interconnected preventing the proper balance of intracellular forces with respect to extracellular adhesion pattern. Therefore, the structural and mechanical interpretations of the adhesive extracellular environment were defective when the crosslinking of actin filaments was reduced.

**Figure 9:**
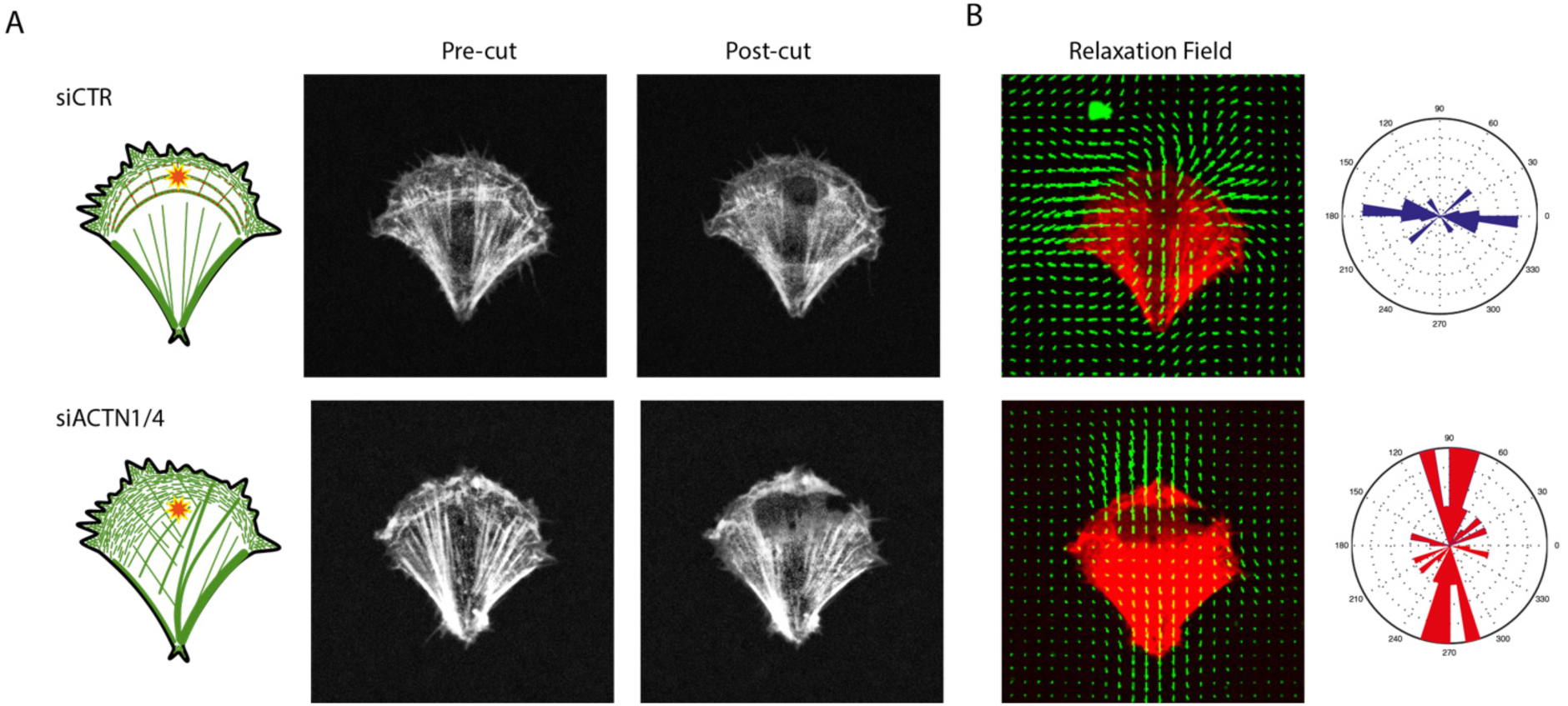
Disruption of the transmission of intracellular tensional forces upon a-actinin depletion. RPE1 cells expressing LifeAct-GFP were seeded on crossbow-shaped micropattern on compliant substrate. (A) Schemes show the localization of nano-ablated regions on control (siCTR; upper row) and a-actinin depleted cells (siACTN1/4; bottom row). Images show actin network before (left) and after (centre) nano-ablation. (B) Images on show the relaxation of the traction force field upon ablation. Graphs show the angular distribution of the principal direction of the relaxation in control (siCTR, n=15) and a-actinin depleted cells (siACTN1/4, n=27).

### Defective integration of spatial cues in α-actinin depleted cells

Our results showed that α-actinin depleted cells were defective in the establishment of balanced actin-network architecture with respect to extracellular cues. We further hypothesized that α-actinin depleted cells are defective in the sensation and adaptation of their architecture in response to local changes in their environment. The absence of large scale connectivity between different cell parts in alpha-actinin depleted cells may impair their ability to integrate local changes in their global internal architecture. To examine this hypothesis, we designed an experiment in which the responses of cells were measured following exposure to anisotropic cues in real time.

The anisotropic cues to plated cells were provided by laser-based micropatterning of additional adhesion sites in situ and in real time (Vignaud et al., 2012b). Two adhesion areas with different distinct distance between adhesion spots were positioned on each side of the symmetry axis (Figure 10A, movie S5). The cell outline was monitored for four hours following the live-patterning of the two distinct adhesive areas (Figure 10B). In control cells and after 1 hour, the protrusions tended to be orientated towards the denser of the two new adhesion sites (Figure 10C). By contrast, in α-actinin depleted cells, the protrusions were orientated randomly with respect to the densities of the adhesion sites (Figure 10C), suggesting that alpha-actinin is a necessary part of the cell’s adaptation to the complexities of the extracellular environment through its role in the integration of diverse mechanical signals from that environment.

**Figure 10:**
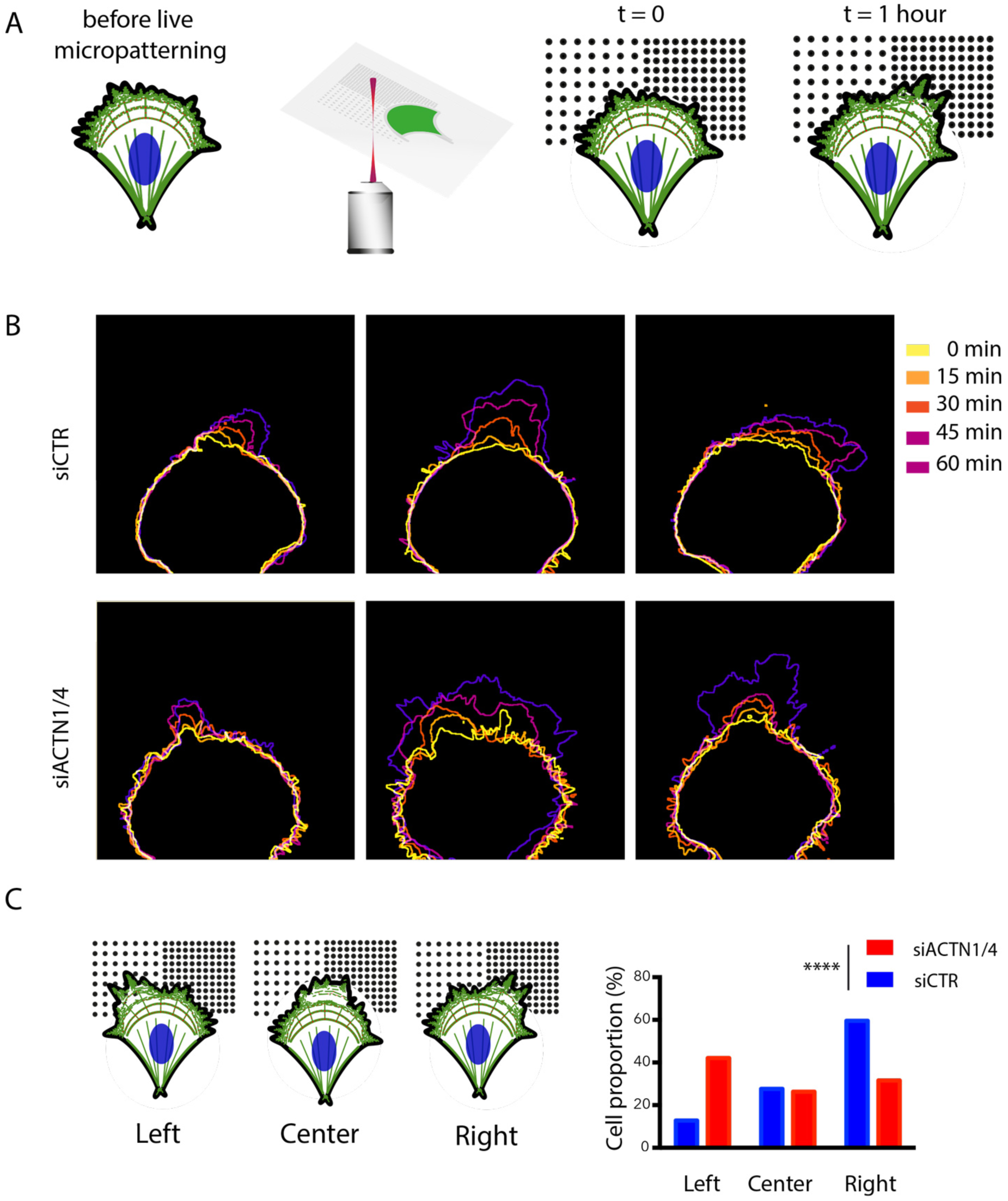
Defective physical integration of external signals upon α-actinin depletion. (A) RPE1 cells were seeded on a crossbow-shaped micropattern (left). A pulsed UV-laser was used to remove anti-fouling coating next to the live cell (centre). Two adhesive areas of different densities were added in front of the cell plated on the crossbow micropattern. The cell was monitored for four hours after the addition of the new patterns. (right). (B) Images show the contour of control (siCTR; upper row) and a-actinin depleted cells (siACTN1/4; bottom row) as cells extend over the new micropatterned adhesion sites. (C) Cell shape was scored one hour after the onset of cell movement. Graphs show the frequency of cell movement toward the left, the centre or the right. Control cells displayed a preferential motion toward the denser adhesive region whereas a-actinin depleted cells could not distinguish the two (siCTR, n=47; siACTN1/4, n=114). Statistical significance was assessed by Chi-square test (p <0.0001).

## Discussion

Our results provide a quantitative description of the remarkable ability of the cell cytoskeleton to self-organize into a contractile architecture that harmonizes with the symmetries and anisotropies of the extracellular environment. The alignment by the retrograde flow of actin filaments into radial fibers attached to cell adhesions and the interconnections of those fibers regulate the transmission of intracellular forces in a cell-wide structure, thus resulting in the geometry of the entire cellular architecture being dictated by the spatial distribution of cell adhesions to the extracellular matrix (Figure 11). Our results revealed that α-actinin, which ensures the connection of transverse arcs with radial fibres and thereby with cell adhesions, plays a key role in ensuring the correct functioning of this mechanism.

**Figure 11:**
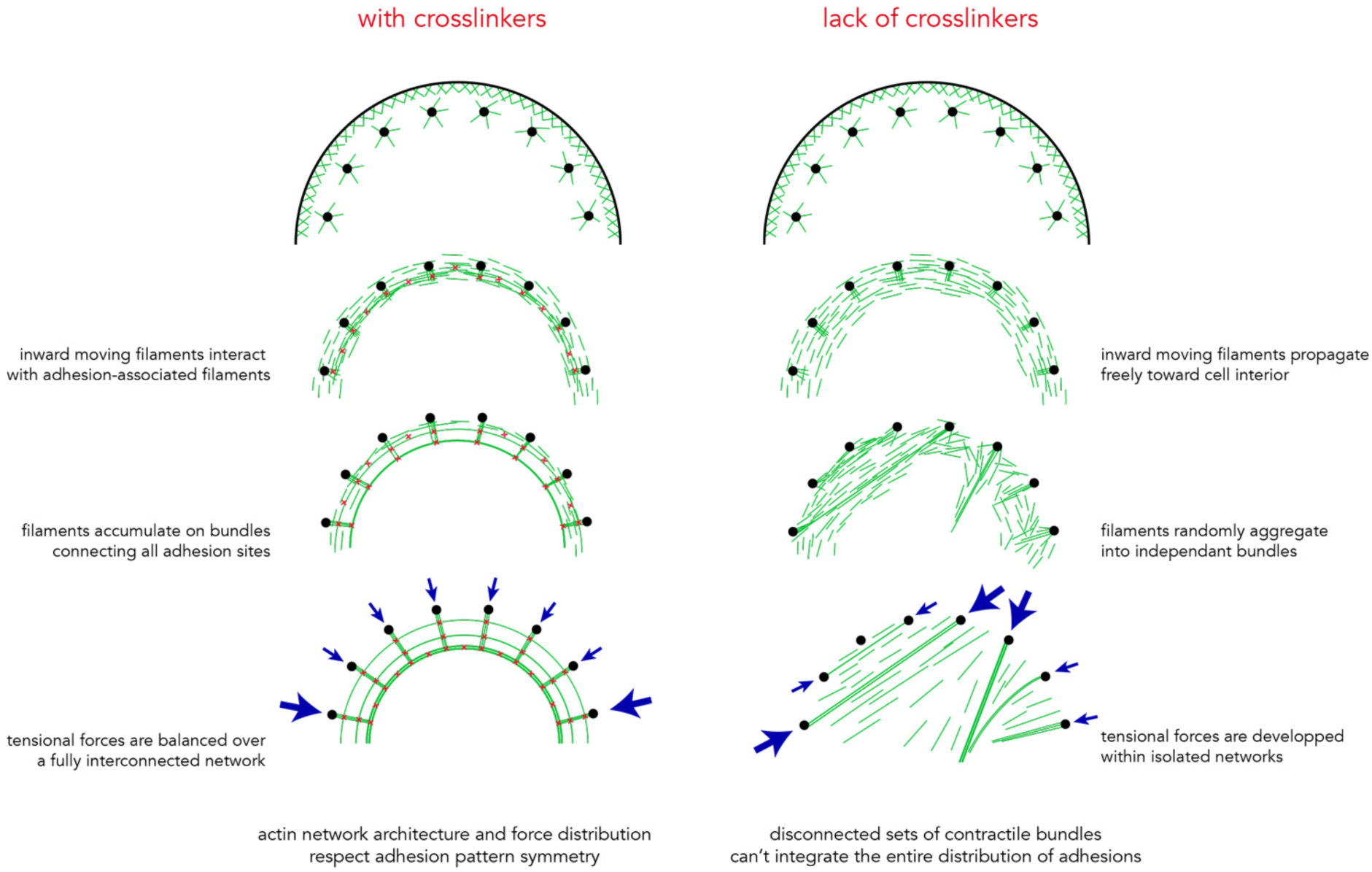
Actin-network dynamics in the cell above the curved adhesive edge in presence or absence of a-actinin. The schemes show the speculative journey of actin filaments from their nucleation (top) to their incorporation into contractile networks (bottom) in control (left) and a-actinin depleted cells (right).

Actin retrograde flow contributes to the establishment of cell polarity (Maiuri et al., 2015). Thus, biased flow might account for biased polarity and the cells inability to properly adapt its architecture to external cues. Furthermore, alpha-actinin accumulation at the transition between the lamellipodium and the lamella, contributes to the local assembly of transverse arcs which further serves as structural supports to the formation of protruding networks that push forward the front edge of migrating cells (Dubin-thaler et al., 2004; Burnette et al., 2011). The role of alpha-actinin in the assembly of these transverse arcs is thus consistent with our observation that cells lacking alpha-actinin are unable to push forward their front extension in the proper direction and a previous report showing that fascin is essential for haptotaxis in fibroblasts (Johnson et al., 2015).

As previously described we found that alpha-actinin knock-down increases the magnitude of traction forces (Oakes et al., 2012). Alpha-actinin has numerous binding partners indicative of functions that extend beyond bundling of actin filaments (Otey and Carpen, 2004; Sjöblom et al., 2008). Notably, α-actinin localizes to cell adhesions where it promotes their maturation (Roca-Cusachs et al., 2013; Meacci et al., 2016). In addition, α-actinin modulates actin dynamics and contractility by competing with other actin binding proteins. For instance α-actinin competes with tropomyosins, which protects actin filaments from cofilin induced severing and activate myosinII (Gateva et al., 2017). This competition determines the rate of actin turn-over and thereby the assembly of stress fiber formation and the regulation of cell motility (Kemp and Brieher, 2018). We also found that alpha-actinin knock-down perturbs the orientation of contractile forces. The two processes may be directly related since high local forces pull on adjacent filaments; their reorientation contributes to reinforce the local contraction but also perturb the orientation of the entire network in a mecano-cumulative effect (Luo et al., 2013; Schiffhauer et al., 2016).

The central role of alpha-actinin in cell architecture, mechanics and mobility may explain cytoskeletal phenotypes observed in certain diseases associated with α-actinin mutations or altered expression levels (Foley and Young, 2014; Murphy and Young, 2015). The balance between ACTN1 and ACTN4 expression levels can lead to different forms of cancer (Sen et al., 2009). A mutant form of ACTN4 which shows a higher affinity to actin as compared to the wild-type isoform, or an increase of ACTN4 concentration, can both lead to focal segmental glomerulosclerosis and renal failure (Feng et al., 2018; Ward et al., 2008). More surprisingly, the same phenotype could be observed in ACTN4 knock-out mice and in humans arboring ACTN4 mutation leading to the expression of an unstable form of the protein (Kos et al., 2003; Bartram et al., 2016). This apparent contradiction raises again the possibility that there is an optimal actinin expression level to ensure connectivity and contraction and proper regulation of tissue morphology (Ennomani et al., 2016; Falkenberg et al., 2017).

The rules describing the physical regulation of cell size and shape are becoming better understood (Marshall, 2015; Paluch and Heisenberg, 2009; Barnhart et al., 2011; Albert and Schwarz, 2014). However, despite a wealth of information about symmetry breaking (Pohl et al., 2015), the rules describing how cell symmetry is regulated are less clear. Cell symetry is a signature of how geometrical and mechanical consistency is maintained within a cell. Marine organisms, which can be considered to be devoid of spatial boundary conditions, display shapes with high levels of symmetry (Sardet, 2015). Recently, jelly-fish symmetry has been shown to be dependent on its contractility (Abrams et al., 2015). At the single-cell level, contractility, mediated by myosin II, was shown to prevent cells from fragmenting (Cai et al., 2010), and to maintain cytoskeletal coherence. Our results suggest that cytoskeletal coherence would also require actin-network crosslinking to ensure that tensional forces are distributed throughout the entire actin network including cell adhesions. In support of this idea, we have shown elsewhere that the force generated by myosins is not sufficient to support actin network contractility, but also requires a precise control of actin-network connectivity (Ennomani et al., 2016). Interestingly, α-actinin, unlike other actin filament crosslinkers, is involved in two key functions (i.e. network connectivity and cell-adhesion maturation) (Peterson et al., 2004; Hotulainen and Lappalainen, 2006; Roca-Cusachs et al., 2013). Hence α-actinin could be viewed as a master integrator of the intracellular constraints created by tensional forces, which ensures cytoskeletal coherence and the alignment of the cytoskeletal architecture with the geometry of extracellular adhesive cues.

## Materials and Methods

### *In-vitro* experiments

Detailed procedure can be found elsewhere (Ennomani et al., 2016). In brief, the Wiskott-Aldrich syndrome pWA domain was adsorbed on the micropatterns and thus promoted actin polymerization upon actin-mix addition. The presence of double headed HMM-myosin-VI was used to induce network contraction (Reymann et al., 2012). The actin mix consisted of 2 µM actin monomers (7% labeled with Alexa 568), 6 µM profilin, 100 nMArp2/3 complex, and 16 nM of HMM-myosin VI (GFP labeled). Proteins were diluted in freshly prepared buffer (15 mM imidazole-HCl (pH 7.8), 0.6 mM ATP, 55 mM DTT, 1 mM EGTA, 75 mM KCl, 3.5 mM MgCl2, 1.5 mg/ml glucose, 10 µg/ml catalase, 50 µg/ml glucose oxidase, and 0.25% w/v methylcellulose). ATP regenerating system was added (2 mM MgATP, 2 mM phosphoenolpyruvate, 2,000 U/ml pyruvate kinase). Network contraction was manipulated by presence (20 nM) or absence of α-actinin 4.

Ring contraction was followed by time-lapse microscopy at one image per minute on an OlympusBX61 microscope with a 40X dry objective (NA = 0.75), a Marzhauser motorized stage and a CoolSnapHQ2 camera (Roper scientific). The system was controlled via Metamorph (Molecular Devices).

### Cell lines

Human telomerase-immortalized, retinal-pigmented epithelial cells (RPE1; Clontech) and RPE1 expressing LifeAct-GFP (Vignaud et al., 2012b) were grown in a humidified incubator at 37°C and 5% CO_2_ in DMEM/F12 medium supplemented with 10% fetal bovine serum and 1% penicillin/streptomycin. All cell culture products were purchased from GIBCO/Life technologies.

### Small-interfering RNA treatment

RPE1 cells were transfected with siRNAs (Qiagen) using lipofectamine RNAi Max transfection reagent (Life Technologies) at a final concentration of 10 nM following the manufacturer’s protocol. Strand sequences were: siACTN4 : 5’-GCAGCAUCGUGGACUACAATT-3’; siACTN1 : 5’-GCACCAUCAUGGACCAUUATT-3’. siRNA specificity has been assessed by two approaches. 1) Redundancy check. We used a second set of siRNA’s directed against ACTN1 and ACTN4 respectively (siACTN1/4 – set02). Strand sequences were as follows: siACTN4 5’-GACACAUAUCGCAGGGAATT-3’ and siACTN1 5’-CACCAUGCAUGCCAUGCAATT-3’. siRNA efficiency has been quantified by western-blot and morphological characterisation after plating on crossbow patterns (Figure S2); and 2) Rescue experiment. Cells were first transfected with siRNA, followed by the transfection of ACTN1-GFP and ACTN4-GFP which are insensitive to the siRNA’s. Transfection was performed with X-tremeGENE HP DNA transfection Reagent (Sigmaaldrich) following manufacturers guidelines. Next cells have been sorted (FACSMELODY – BD Bioscience) so as to isolate two populations R- (cells which do not harbour the rescue constructs) and R+ (GFP positif). Efficiency of the approach has been assessed by western-blot and morphological analysis after plating on crossbow patterns (Figure S2 and Figure 2). For morphological analysis only cells with consistent actinin localisation were taken into account in order to avoid over-expression artifacts.

### Antibodies and cytoskeletal labelling agents

Primary antibodies were obtained from the following sources and used at the following dilutions: rabbit anti-GAPDH (Santa Cruz #25778; 1/2000 for western blot), rabbit anti-phospho myosin light chain 2 (Ser19) (Cell Signaling Technology #3671; 1/50 for Iimmunofluorescence), rabbit anti-ACTN1 (Sigma #HPA006035) 1/500 WB 1/100 IF, mouse anti-ACTN4 (abcam ab59458 1/1000 WB, 1/100 IF), rabbit anti-myosin IIa (Cell Signaling Technology #3403; 1/100 for IF), rabbit anti-myosin IIb (Cell Signaling Technology #3404; 1/200 for IF). Alexa fluorophore-conjugated phalloidin (Molecular Probes) was resuspended in methanol and diluted 1/500 in PBS for immunostaining. Alexa fluorophore-conjugated secondary antibodies (Molecular Probes) were diluted 1/1000.

### Immunostaining

Cells were fixed in 4% paraformaldehyde in cytoskeleton buffer pH 6,1 for 15 min at room temperature. They were then washed twice with PBS and incubated in quenching agent 0,1 ammonium chloride in PBS for 10 min. Cells were then permeabilized in 0.1% Triton X-100 in PBS for 3 min. For phospho-myosin immunostaining, cells were prepermeabilized for 15 seconds with 0.1% Triton X-100 in cytoskeleton buffer prior to paraformaldehyde fixation for 15 min. Paraformaldehyde autofluorescence was quenched by ammonium chloride incubation.

For all conditions, after fixation, the cells were washed then blocked with 1.5% bovine serum albumin (BSA) for 30 minutes. The cells were stained with primary antibodies diluted in PBS with 1.5% BSA for 1 hour, followed by extensive washing with PBS and staining with secondary antibodies diluted in PBS with 1.5% BSA for 30 min. After washing, Alexa fluorophore-conjugated phalloidin was incubated 20 min. The cells were washed with PBS and the DNA labelled with 0.2 µg/ml DAPI (Sigma) for 10 minutes. After washing the cells with water, the coverslips were air-dried and mounted onto slides using Mowiol.

### Image acquisition

Images of fixed cells were acquired using either an upright microscope (BX61; Olympus) with 100X (1.4 N.A) oil immersion objective mounted on a piezo ceramic (Physics Instruments), controlled with MetaMorph, or a Leica-TCS-SP2 confocal microscope with a 63X (0.9 N.A) oil immersion objective.

### Western Blotting

RPE1 cells were lysed at 4°C in RIPA buffer (Thermo Scientific) supplemented with 1x protease inhibitor cocktail (Roche). Supernatants were collected and loaded on SDS polyacrylamide gel electrophoresis and transferred onto nitrocellulose membrane using a semi-dry Western blotting apparatus (BioRad). The membranes were blocked with PBS containing 5% non-fat milk for 1 hour at room temperature. After blocking the membranes were probed with primary antibodies overnight at 4°C. The membranes were washed four times with blocking buffer before adding HRP-conjugated secondary antibodies (Life Technologies), diluted as recommended in blocking buffer, and incubating for 30 min at ambient temperature. After washing three times with PBS containing 0.1% Tween-20 (Sigma) the membranes were developed using ECL reagent (Life Technologies) and imaged on the ChemiDoc system (BioRad) or by exposing to scientific imaging film (Kodak).

### Glass Patterning

Detailed procedure has been already described elsewhere (Azioune et al., 2010). In brief, glass coverslips were spin-coated (30 sec, 3000 rpm) with adhesion promoter Ti-Prime (MicroChemicals), then cured for 5 min at 120°C and further spin-coated (30 sec., 1000 rpm) with 1% polystyrene (Sigma) in toluene (Sigma). Coated coverslips were next oxidized by oxygen plasma (FEMTO, Diener Electronics) (10 sec, 30 W) and incubated for 30 min. with 0.1 mg/ml PLL-g-PEG (PLL20K-G35-PEG2K, JenKem) in 10mM HEPES pH 7.4. Dried coverslips where then exposed to deep-UV (UVO cleaner, Jelight) through a photomask (Toppan) for 4 min. After UV treatment, coverslips were incubated with 10 µg/ml fibronectin (Sigma) and 10 µg/ml Alexa Fluor 546 fibrinogen conjugate (Invitrogen) in PBS for 30 min.

### Soft patterning

Detailed procedure has been already described elsewhere (Vignaud et al., 2014). Quartz photomask was oxidized through oxygenplasma (Femto, Diener Electronics) for 3 min at 100 W before incubation with 0.1 mg/ml PLL-g-PEG in 10mM HEPES, pH 7.4, for 30 min. After drying, uncoated photomask side was exposed to deep-UV for 5 min. Next, the PLL-g-PEG coated side was incubated with 10 µg/ml fibronectin (Sigma) and 10 µg/ml Alexa Fluor 647 fibrinogen conjugate (Invitrogen) in 100mM sodium bicarbonate buffer, pH=8.4, for 30 min. Acrylamide (8%) and bis-acrylamide solution (0.264%) (Sigma) was degassed for 30 min, mixed with passivated fluorescent beads (Invitrogen) by sonication before addition of APS and TEMED. 25 µl of that solution are applied on the micropatterned photomask, covered with a silanized coverslip and allow to polymerize for 30 min. Protocols for bead passivation and glass silanization can be found elsewhere (Tseng et al., 2012). The gel was allowed to swell in 100mM sodium bicarbonate buffer and gently removed. Coverslip were rinsed with PBS before cell plating.

The Young-modulus of the gels was measured around 35kPa by atomic force microscopy.

### Traction force microscopy

Imaging was performed with a confocal spinning disk system consisting of an EclipseTi-E Nikon inverted microscope, equipped with a CSUX1-A1 Yokogawa confocal head, an Evolve EMCCD camera (Roper Scientific, Princeton Instruments) and a Nikon CFI Plan-APO VC 60X, 1.4 N.A, oil-immersion objective, interfaced with MetaMorph (Universal Imaging).

Displacement fields were obtained from bead images before and after removal of cells by trypsin treatment. Bead images were first aligned to correct for experimental drift. The displacement field was calculated by particle imaging velocimetry (PIV) on the basis of normalized cross-correlation following an iterative scheme. Final grid size was 0.267 µm X 0.267 µm. Erroneous vectors where discarded on the basis of a low correlation value and replaced by the median value of the neighbouring vectors. The traction-force field was subsequently reconstructed by Fourier Transform Traction Cytometry, with a regularization parameter set to 2×10^−10^. Detailed procedures and code are freely available (Martiel et al., 2015).

Force dipole analysis was performed as described previously (Butler et al., 2002; Mandal et al., 2014) using a custom written Matlab code.

### Laser dissection

UV-laser-based nano-ablation was performed on the spinning disk system using the iLas2 device (Roper Scientific, France). iLas2 is a dual axis galvanometer based optical scanner that focuses the laser beam on the sample. The system includes a telescope to adjust laser focus and a motorized polarizer to control beam power. We used a passively Q-switched laser (STV-E, TeamPotonics, France) producing 300 picoseconds pulses at 355 nm). Laser displacement, exposure time and repetition rate were controlled via ILas software interfaced with MetaMorph (Universal Imaging Corporation). Laser ablation and subsequent imaging was performed with a 100 X CFI S Fluor oil objective (MRH02900, Nikon). The ablation region was exposed for 12 msec at a repetition rate of 7000 Hz. Pulse energy was set to 300 nJ before objective by adjusting the polarizer.

### Live laser patterning

RPE1-LA-GFP cells were seeded on patterned coverslips without prior protein coating and allowed to attach for 3 hours. Live laser patterning was performed as described previously (Vignaud et al., 2012b). In brief, patterning was performed on the aforementioned spinning-disk system (see Section Traction Force Microscopy), combined with iLas2 (Roper Scientific, France) and a 100X CFI S Fluor oil objective (MRH02900, Nikon). We used a passively Q-switched laser with a 355 nm wavelength. Patterns were designed in Illustrator (Adobe) and imported into iLas software. Typical patterns consist of arrays of 1 µm diameter spots, with a pitch of 3 µm (high spot density) or 4 µm (low spot density).

### Transverse arcs tracking

RPE1-LA-GFP cells were seeded on patterned coverslips and allowed to spread for 3 hours. Time-lapse images were acquired on spinning-disk system with a Nikon CFI PlanApo 60X oil objective (N.A. 1.4). Over a period of 20 min, we acquired, every minute, a Z-stack consisting of five planes covering 5 µm of the basal plane of the cell. Illumination was set so as to have highest dynamical range at the leading edge of the cell. Resulting stacks were analysed by the “Analyze Tubeness” function bundled with the freely available software FIJI (https://fiji.sc/). Particle image velocymetry (see Section Traction Force Microscopy) was performed on maximum projection of each stack with the freely available software PIVLab (Thielicke and Stamhuis, 2014). PIVlab was employed by performing a double cross correlation analysis, with an interrogation area of 30 pixels with a step of 15 pixels. We produced average flow fields giving access to mean flow speed and orientation.

### Image analysis

#### Directionality

Each cell was divided in two equal regions of interest (ROI) following the patterns symmetry axis. Actin-fibre orientation was assessed with two freely available imageJ plugins (https://imagej.nih.gov/ij/): Directionality, as implemented by J-Y Tinevez (http://imagej.net/Directionality), and OrientationJ, as implemented by D. Sage (http://bigwww.epfl.ch/demo/orientation/).

Orientation distribution were compared by Kolmogorov-Smirnov test, where D is the difference between fibre orientation distribution between each part of the cell. Both approaches gave similar results.

#### Fluorescence measurement

MyosinIIA, IIB and pMLC signals were analyzed with the freely available software imageJ (https://imagej.nih.gov/ij/). Each cell was divided in two equal ROIs, following the patterns symmetry axis. Fluorescence intensity was measured in each ROI. The ratio of fluorescence intensity between each region of interest describes to which extend the signal distribution is symmetric with respect to the patterns symmetry axis.

#### Connectivity analysis

Actin cytoskeleton was segmented following two strategies based on ridge detection. First approach was based on ridge detection with SteerableJ an imageJ plugin implemented by François Aguet (http://bigwww.epfl.ch/demo/steerable/download.html) (Jacob and Unser, 2004) and second approach used Ridge_Detection as implemented by Wagner and Hiner (http://imagej.net/Ridge_Detection)(Steger, 1998). Both approaches produced similar structures. Next, binary image was skeletonized and analyzed using imageJ. Among different parameters, and statistics, which describe the skeleton architecture number of junctions and skeleton length allow to quantify the network connectivity. Skeleton length refers to the biggest skeleton in the considered cell and is the product between number of branches and average branch length as given by the ImageJ function “Analyze skeleton”.

#### Nucleus positioning

DAPI stained nuclei were first LoG filtered. Images were thresholded with automatic Otsu approach and segmented. Image processing was performed in imageJ. Nuclei coordinates were approximated to the centre of mass of the segmented nucleus. These coordinates were measured relatively to the geometric centre of the micropattern.

## Supplementary Figures

**Figure S1:**
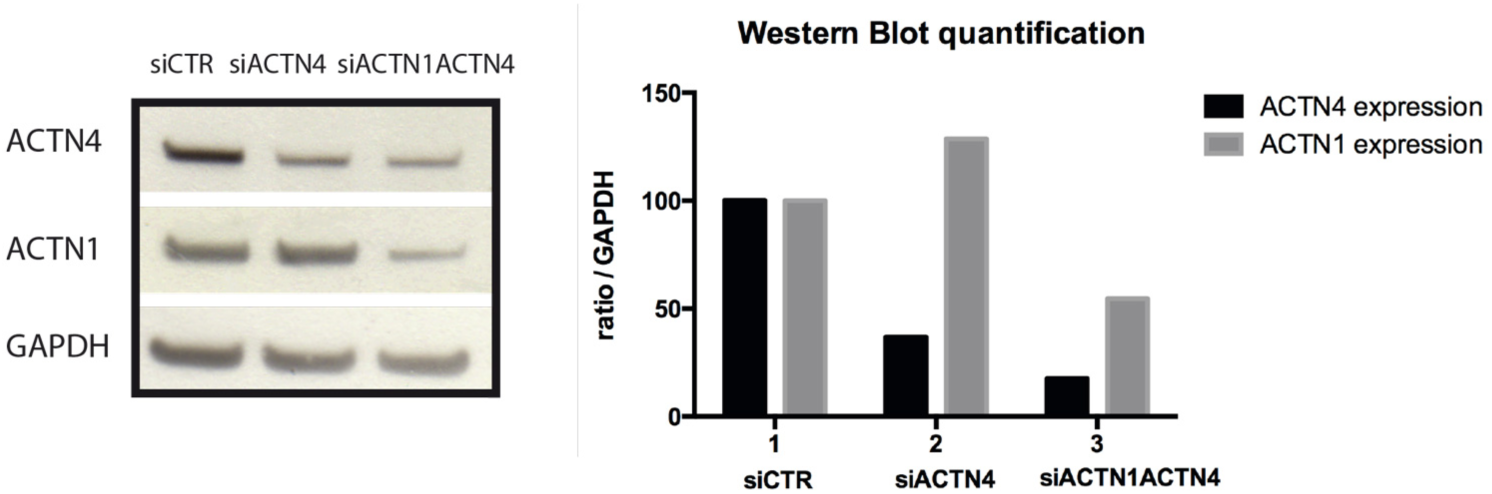
siRNA efficiency. Silencing-RNA efficiency was assessed by western-blotting. GAPDH expression was used as reference. Quantification of expression levels was calculated as protein expression level to GAPDH expression level ratio. siACTN4 treatment alone induced overexpression of siACTN1. Using siACTN4 and siACTN1 in combination, significantly decreased expression levels for both proteins.

**Figure S2:**
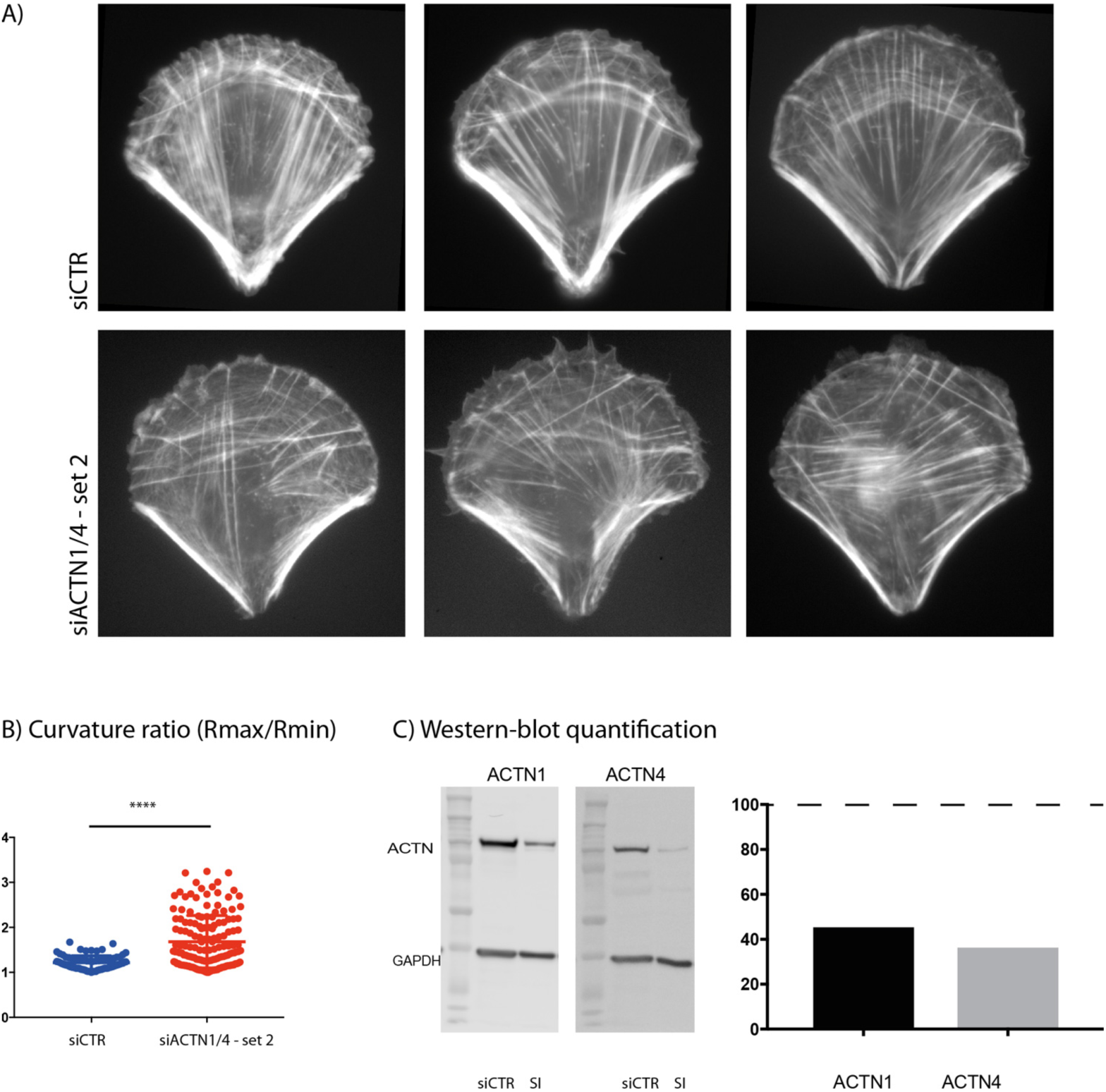
siRNA specificity / Redundancy check. A) Typical images of RPE1-LA-GFP cells seeded on crossbow patterns after either treatment with control siRNA or siACTN1/4-set2. B) Morphological characterisation of the cell. Radius of curvature for each flanking stress fiber has been measured. The plot depicts the ratio of the biggest to the smallest radius for control cells (siCTR; mean = 1,232; n=79) and siACTN1/4-set2 (mean = 1,861; n=162). (Statistical significance has been assessed by Mann-Whithney test (p<0,0001). C) Western-blot analysis and expression level quantification. Control levels have been set to 100% (dotted line) after normalization with GAPDH expression level. ACTN1 and ACTN4 levels are given relative to control. Overall actin architecture, cellular morphology and protein expression levels are in accordance with the results obtained with siACTN1/4. (see figure 2, 3 and S1).

**Figure S3:**
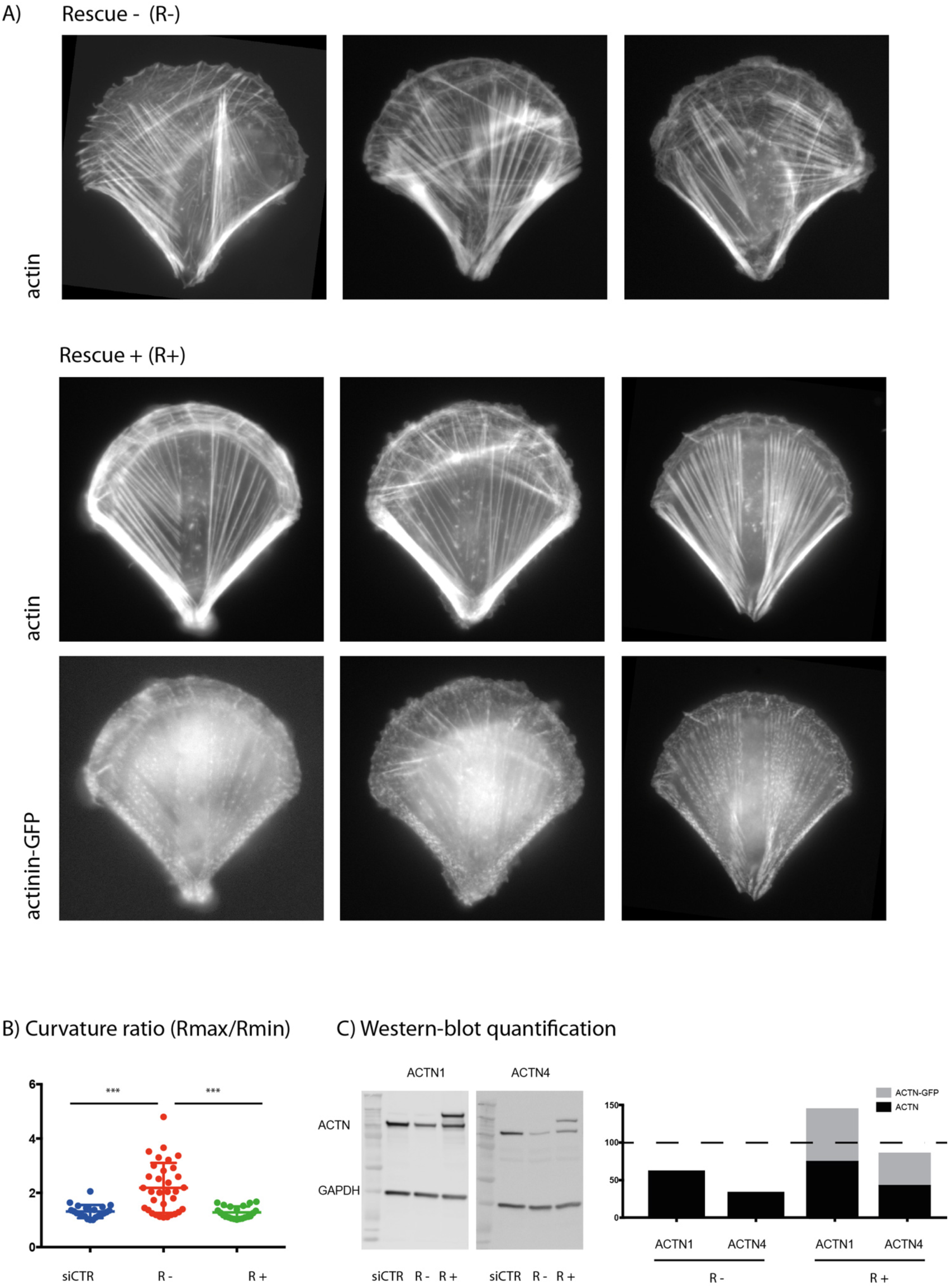
siRNA specificity / Rescue. A) Typical images of cells treated with siRNA1/4, subsequently transfected with ACTN1-GFP and ACTN4-GFP. Cells have been sorted so as to isolate two populations Rescue – (R-) which didn’t incorporated any of the ACTN constructs, and Rescue+ (R+) the GFP positive population. B) Morphological characterization has been performed by measuring the ratio of curvature. Curvature radius has been measured for each flanking stress fiber and reported as the ratio of the biggest (Rmax) to the smallest (Rmin). siCTR (mean=1,311; n=22). R-(mean=2,181; n=33). R+ (mean=1,275; n=22). Multiple comparison has been performed by a Kruskal-Wallis test (siCTR vs R-, p=0,0004; siCTR vs R+, p>0,9999; R-vs R+, p = 0,0001). C) Western-blot analysis and expression level quantification. Control levels have been set to 100% (dotted line) after normalization with GAPDH expression level. ACTN1 and ACTN4 levels are given relative to control. This ensemble measurement shows the efficiency of the siRNA approach as R-cells show significantly decreased expression levels for ACTN1 and ACTN4. Expression levels for R+ cells sum up endogenous ACTN, wich was efficiently down-regulated by siRNA, and ACTN-GFP expression level. We therefore notice a global over-expression for ACTN1 (144 %) and an almost basal expression level for ACTN4 (85%).

## Supplementary Movies

**Movie S1:** In vitro reconstitution of contractile actin rings.

This movie shows the contraction of networks of actin filaments (red) assembled on a 60-micron-wide 18-dot micropattern of pWA upon the addition of double headed GFP-myosin-VI (green). The white circle and the white cross mark the initial shape of the ring and its centre. Actinin was added only on the right movie at a final concentration of 20 nM.

**Movie S2:** formation and propagation of actin arcs.

This movie shows the dynamic of actin network (Lifeact-GFP) in RPE1 cells plated on crossbow-shaped micropattern. Cells were treated with a control siRNA (siCTR, left) or siRNAs against alpha-actinin 1 and 4 (siACTN, right). Images were taken every minute. The inset on the right is a magnified view of the upper-right part of the cell. Actin arcs formed in retracting protrusions and concatenated in a continuous cable lining the cell front as they contracted and moved inward.

**Movie S3:** Retrograde actin flow.

This movie shows the dynamic of actin network (Lifeact-GFP) in RPE1 cells plated on crossbow-shaped micropattern. Cells were treated with a control siRNA (siCTR, left) or siRNAs against alpha-actinin 1 and 4 (siACTN, right). Images were taken every 30 seconds. In each pannel, the left movie shows raw images and the right movies the segmented images. The upper inset is a magnified view of the upper part of the cell. Red arrows show the local orientation and magnitude of actin structure displacements.

**Movie S4**: Traction force relaxation upon central fiber ablation.

This movie shows the relaxation of the traction force field of RPE1 cells plated on crossbow-shaped micropattern on poly-acrylamide hydrogels. Cells were expressing Lifeact-GFP in order to visualize the actin network (gray). Cells were treated with a control siRNA (siCTR, top) or siRNAs against alpha-actinin 1 and 4 (siACTN, bottom). Images were taken every 5 seconds. Green arrows show the displacement field of the beads embedded in the acrylamide gels between two consecutive images. The red arrows on the second image show the ablated region. As a consequence, the relaxation field is much higher on this image and differentially orientated in control cells and those in which alpha-actinin has been down regulated.

**Movie S5:** Cell motion in response to differential ECM densities.

This movie shows RPE1 cells expressing Lifeact-GFP and moving out of crossbow-shaped micropattern in response to substrate live micropatterning. High (on the right) and low densities (on the left) of ECM spots were added in front of spatially confined cells, offering them the possibility to move on these newly presented regions. Cells were treated with a control siRNA (siCTR, top) or siRNAs against alpha-actinin 1 and 4 (siACTN, bottom). Images were taken every 10 minutes.

## References

Abrams, M.J., T. Basinger, W. Yuan, C.-L. Guo, and L. Goentoro. 2015. Self-repairing symmetry in jellyfish through mechanically driven reorganization. Proc. Natl. Acad. Sci. 112:E3365–E3373. doi:10.1073/pnas.1502497112.

Abu Shah, E., and K. Keren. 2014. Symmetry breaking in reconstituted actin cortices. Elife. 3:1–15. doi:10.7554/eLife.01433.

Albert, P.J., and U.S. Schwarz. 2014. Dynamics of Cell Shape and Forces on Micropatterned Substrates Predicted by a Cellular Potts Model. Biophys. J. 106:2340–2352. doi:10.1016/j.bpj.2014.04.036.

Ashdown, G.W., G.L. Burn, D.J. Williamson, E. Pandžić, R. Peters, M. Holden, H. Ewers, L. Shao, P.W. Wiseman, and D.M. Owen. 2017. Live-Cell Super-resolution Reveals F-Actin and Plasma Membrane Dynamics at the T Cell Synapse. Biophys. J. 112:1703–1713. doi:10.1016/j.bpj.2017.01.038.

Azioune, A., N. Carpi, Q. Tseng, M. Théry, and M. Piel. 2010. Protein micropatterns: A direct printing protocol using deep UVs. Methods Cell Biol. 97:133–46. doi:10.1016/S0091-679X(10)97008-8.

Barnhart, E.L., K.-C. Lee, K. Keren, A. Mogilner, and J. a Theriot. 2011. An Adhesion-Dependent Switch between Mechanisms That Determine Motile Cell Shape. PLoS Biol. 9:e1001059. doi:10.1371/journal.pbio.1001059.

Bartram, M.P., S. Habbig, C. Pahmeyer, M. Höhne, L.T. Weber, H. Thiele, J. Altmüller, N. Kottoor, A. Wenzel, M. Krueger, B. Schermer, T. Benzing, M.M. Rinschen, and B.B. Beck. 2016. Three-layered proteomic characterization of a novel *ACTN4* mutation unravels its pathogenic potential in FSGS. Hum. Mol. Genet. 25:1152–1164. doi:10.1093/hmg/ddv638.

Belmonte, J., M. Leptin, and F. Nedelec. 2017. A Theory That Predicts Behaviors Of Disordered Cytoskeletal Networks. bioRxiv.

Blanchoin, L., R. Boujemaa-Paterski, C. Sykes, and J. Plastino. 2014. Actin dynamics, architecture, and mechanics in cell motility. Physiol. Rev. 94:235–63. doi:10.1152/physrev.00018.2013.

Burnette, D.T., S. Manley, P. Sengupta, R. Sougrat, M.W. Davidson, B. Kachar, and J. Lippincott-schwartz. 2011. A role for actin arcs in the leading-edge advance of migrating cells. Nat. Cell Biol. 13. doi:10.1038/ncb2205.

Butler, J.P., I.M. Tolić-Nørrelykke, B. Fabry, and J.J. Fredberg. 2002. Traction fields, moments, and strain energy that cells exert on their surroundings. Am. J. Physiol. Cell Physiol. 282:C595–605. doi:10.1152/ajpcell.00270.2001.

Cai, Y., O. Rossier, N.C. Gauthier, N. Biais, M.-A. Fardin, X. Zhang, L.W. Miller, B. Ladoux, V.W. Cornish, and M.P. Sheetz. 2010. Cytoskeletal coherence requires myosin-IIA contractility. J. Cell Sci. 123:413–23. doi:10.1242/jcs.058297.

Chang, C., and S. Kumar. 2013. Vinculin tension distributions of individual stress fibers within cell-matrix adhesions. J. Cell Sci. 126:3021–30. doi:10.1242/jcs.119032.

Chau, A.H., J.M. Walter, J. Gerardin, C. Tang, and W.A. Lim. 2012. Designing synthetic regulatory networks capable of self-organizing cell polarization. Cell. 151:320–332. doi:10.1016/j.cell.2012.08.040.

Ciobanasu, C., B. Faivre, and C. Le Clainche. 2014. Actomyosin-dependent formation of the mechanosensitive talin-vinculin complex reinforces actin anchoring. Nat. Commun. 5:3095. doi:10.1038/ncomms4095.

Dubin-thaler, B.J., N. Kieffer, A.R. Bresnick, and M.P. Sheetz. 2004. Periodic Lamellipodial Contractions Correlate with Rearward Actin Waves. 116:431–443.

Ennomani, H., G. Letort, C. Guérin, J.-L. Martiel, W. Cao, F. Nédélec, E.M. De La Cruz, M. Théry, and L. Blanchoin. 2016b. Architecture and Connectivity Govern Actin Network Contractility. Curr. Biol. 26:616–26. doi:10.1016/j.cub.2015.12.069.

Falkenberg, C. V, E.U. Azeloglu, M. Stothers, T.J. Deerinck, Y. Chen, J.C. He, M.H. Ellisman, J.C. Hone, R. Iyengar, and M. Loew. 2017. Fragility of foot process morphology in kidney podocytes arises from chaotic spatial propagation of cytoskeletal instability. 1–21. doi:10.5061/dryad.09d0k.

Feng, D., J. Notbohm, A. Benjamin, S. He, M. Wang, L.-H. Ang, M. Bantawa, M. Bouzid, E. Del Gado, R. Krishnan, and M.R. Pollak. 2018. Disease-causing mutation in α-actinin-4 promotes podocyte detachment through maladaptation to periodic stretch. Proc. Natl. Acad. Sci. U. S. A. 201717870. doi:10.1073/pnas.1717870115.

Fletcher, D. a, and R.D. Mullins. 2010. Cell mechanics and the cytoskeleton. Nature. 463:485–92. doi:10.1038/nature08908.

Foley, K.S., and P.W. Young. 2014. The non-muscle functions of actinins: an update. Biochem. J. 459:1–13. doi:10.1042/BJ20131511.

Gateva, G., E. Kremneva, T. Reindl, T. Kotila, K. Kogan, L. Gressin, P.W. Gunning, D.J. Manstein, A. Michelot, and P. Lappalainen. 2017. Tropomyosin Isoforms Specify Functionally Distinct Actin Filament Populations In Vitro. Curr. Biol. 1–9. doi:10.1016/j.cub.2017.01.018.

Gomes, E.R., S. Jani, and G.G. Gundersen. 2005. Nuclear movement regulated by Cdc42, MRCK, myosin, and actin flow establishes MTOC polarization in migrating cells. Cell. 121:451–63. doi:10.1016/j.cell.2005.02.022.

Hotulainen, P., and P. Lappalainen. 2006. Stress fibers are generated by two distinct actin assembly mechanisms in motile cells. J. Cell Biol. 173:383–94. doi:10.1083/jcb.200511093.

Ingber, D.E. 2003. Tensegrity II. How structural networks influence cellular information processing networks. J. Cell Sci. 116:1397–1408. doi:10.1242/jcs.00360.

Jacob, M., and M. Unser. 2004. Design of steerable filters for feature detection using Canny-like criteria. IEEE Trans. Pattern Anal. Mach. Intell. 26:1007–1019. doi:10.1109/TPAMI.2004.44.

Jensen, M.H., E.J. Morris, and D.A. Weitz. 2015. Mechanics and dynamics of reconstituted cytoskeletal systems. Biochim. Biophys. Acta - Mol. Cell Res. 1853:3038–3042. doi:10.1016/j.bbamcr.2015.06.013.

Jiu, Y., J. Lehtimäki, S. Tojkander, F. Cheng, H. Jäälinoja, X. Liu, M. Varjosalo, J.E. Eriksson, and P. Lappalainen. 2015. Bidirectional Interplay between Vimentin Intermediate Filaments and Contractile Actin Stress Fibers. Cell Rep. 11:1511–1518. doi:10.1016/j.celrep.2015.05.008.

Johnson, H.E., S.J. King, S.B. Asokan, J.D. Rotty, J.E. Bear, and J.M. Haugh. 2015. F-actin bundles direct the initiation and orientation of lamellipodia through adhesion-based signaling. J. Cell Biol. 208:443–455. doi:10.1083/jcb.201406102.

Kemp, J.P., and W.M. Brieher. 2018. The actin filament bundling protein α-actinin-4 actually suppresses actin stress fibers by permitting actin turnover. J. Biol. Chem. 293:14520–14533. doi:10.1074/jbc.RA118.004345.

Kos, C.H., T.C. Le, S. Sinha, J.M. Henderson, S.H. Kim, H. Sugimoto, R. Kalluri, R.E. Gerszten, and M.R. Pollak. 2003. Mice deficient in alpha-actinin-4 have severe glomerular disease. J. Clin. Invest. 111:1683–90. doi:10.1172/JCI17988.

Kovac, B., J.L. Teo, T.P. Mäkelä, and T. Vallenius. 2013. Assembly of non-contractile dorsal stress fibers requires α-actinin-1 and Rac1 in migrating and spreading cells. J. Cell Sci. 126.

Kumar, S., I.Z. Maxwell, A. Heisterkamp, T.R. Polte, T.P. Lele, M. Salanga, E. Mazur, and D.E. Ingber. 2006. Viscoelastic retraction of single living stress fibers and its impact on cell shape, cytoskeletal organization, and extracellular matrix mechanics. Biophys. J. 90:3762–73. doi:10.1529/biophysj.105.071506.

Lee, S., and S. Kumar. 2016. Actomyosin stress fiber mechanosensing in 2D and 3D. F1000Research. 5:2261. doi:10.12688/f1000research.8800.1.

Letort, G., H. Ennomani, L. Gressin, M. Théry, L. Blanchoin, G. Letort, H. Ennomani, L. Gressin, M. Théry, and L. Blanchoin. 2015. Dynamic reorganization of the actin cytoskeleton. F1000Research. 4. doi:10.12688/f1000research.6374.1.

Levayer, R., and T. Lecuit. 2012. Biomechanical regulation of contractility: spatial control and dynamics. Trends Cell Biol. 22:61–81. doi:10.1016/j.tcb.2011.10.001.

Lim, W.A., C.M. Lee, and C. Tang. 2013. Design Principles of Regulatory Networks: Searching for the Molecular Algorithms of the Cell. Mol. Cell. 49:202–212. doi:10.1016/j.molcel.2012.12.020.

Lim, Y., S.-T. Lim, A. Tomar, M.L. Gardel, J.A. Bernard-Trifilo, X.L. Chen, S.A. Uryu, R. Canete-Soler, J. Zhai, H. Lin, W.W. Schlaepfer, P. Nalbant, G. Bokoch, D. Ilic, C.M. Waterman-Storer, and D.D. Schlaepfer. 2008. PyK2 and FAK connections to p190Rho guanine nucleotide exchange factor regulate RhoA activity, focal adhesion formation, and cell motility. J. Cell Biol. 180:187–203. doi:10.1083/jcb.200708194.

Linsmeier, I., S. Banerjee, P.W. Oakes, W. Jung, T. Kim, and M.P. Murrell. 2016. Disordered actomyosin networks are sufficient to produce cooperative and telescopic contractility. Nat. Commun. 7:12615. doi:10.1038/ncomms12615.

Luo, T., K. Mohan, P.A. Iglesias, and D.N. Robinson. 2013. Molecular mechanisms of cellular mechanosensing. Nat. Mater. 12:1064–1071. doi:10.1038/nmat3772.

Maiuri, P., J.-F. Rupprecht, S. Wieser, V. Ruprecht, O. Bénichou, N. Carpi, M. Coppey, S. De Beco, N. Gov, C.-P. Heisenberg, C. Lage Crespo, F. Lautenschlaeger, M. Le Berre, A.-M. Lennon-Dumenil, M. Raab, H.-R. Thiam, M. Piel, M. Sixt, and R. Voituriez. 2015. Actin Flows Mediate a Universal Coupling between Cell Speed and Cell Persistence. Cell. 161:374–386. doi:10.1016/J.CELL.2015.01.056.

Mak, M., M.H. Zaman, R.D. Kamm, and T. Kim. 2016. Interplay of active processes modulates tension and drives phase transition in self-renewing, motor-driven cytoskeletal networks. Nat. Commun. 7:10323. doi:10.1038/ncomms10323.

Mandal, K., I. Wang, E. Vitiello, L.A.C. Orellana, and M. Balland. 2014. Cell dipole behaviour revealed by ECM sub-cellular geometry. Nat. Commun. doi:10.1038/ncomms6749.

Marshall, W.F. 2015. Cell Geometry: How Cells Count and Measure Size. Annu. Rev. Biophys. 49–64. doi:10.1146/annurev-biophys-062215-010905.

Martiel, J.-L., A. Leal, L. Kurzawa, M. Balland, I. Wang, T. Vignaud, Q. Tseng, and M. Théry. 2015. Measurement of cell traction forces with ImageJ. Methods Cell Biol. 125:269–87. doi:10.1016/bs.mcb.2014.10.008.

Meacci, G., H. Wolfenson, S. Liu, M.R. Stachowiak, T. Iskratsch, A. Mathur, S. Ghassemi, N. Gauthier, E. Tabdanov, J. Lohner, A. Gondarenko, A.C. Chander, P. Roca-Cusachs, B. O’Shaughnessy, J. Hone, and M.P. Sheetz. 2016. α-actinin links ECM rigidity sensing contractile units with periodic cell edge retractions. Mol. Biol. Cell. doi:10.1091/mbc.E16-02-0107.

Mukhina, S., Y.-L. Wang, and M. Murata-Hori. 2007. Alpha-actinin is required for tightly regulated remodeling of the actin cortical network during cytokinesis. Dev. Cell. 13:554–65. doi:10.1016/j.devcel.2007.08.003.

Murphy, A.C.H., and P.W. Young. 2015. The actinin family of actin cross-linking proteins - a genetic perspective. Cell Biosci. 5:49. doi:10.1186/s13578-015-0029-7.

Murrell, M.P., and M.L. Gardel. 2012. F-actin buckling coordinates contractility and severing in a biomimetic actomyosin cortex. Proc. Natl. Acad. Sci. 109:20820–20825. doi:10.1073/pnas.1214753109.

Naumanen, P., P. Lappalainen, and P. Hotulainen. 2008. Mechanisms of actin stress fibre assembly. In Journal of Microscopy. 446–454.

Oakes, P.W., Y. Beckham, J. Stricker, and M.L. Gardel. 2012. Tension is required but not sufficient for focal adhesion maturation without a stress fiber template. J. Cell Biol. 196:363–74. doi:10.1083/jcb.201107042.

Otey, C.A., and O. Carpen. 2004. Alpha-actinin revisited: a fresh look at an old player. Cell Motil. Cytoskeleton. 58:104–11. doi:10.1002/cm.20007.

Paluch, E., and C.-P. Heisenberg. 2009. Biology and physics of cell shape changes in development. Curr. Biol. 19:R790–9. doi:10.1016/j.cub.2009.07.029.

Peterson, L.J., Z. Rajfur, A.S. Maddox, C.D. Freel, Y. Chen, M. Edlund, C. Otey, and K. Burridge. 2004. Simultaneous stretching and contraction of stress fibers in vivo. Mol. Biol. Cell. 15:3497–508. doi:10.1091/mbc.E03-09-0696.

Pohl, C., Pohl, and Christian. 2015. Cytoskeletal Symmetry Breaking and Chirality: From Reconstituted Systems to Animal Development. Symmetry (Basel). 7:2062–2107. doi:10.3390/sym7042062.

Reymann, A.-C., R. Boujemaa-Paterski, J.-L. Martiel, C. Guérin, W. Cao, H.F. Chin, E.M. De La Cruz, M. Théry, and L. Blanchoin. 2012. Actin network architecture can determine myosin motor activity. Science. 336:1310–4. doi:10.1126/science.1221708.

Reymann, A., J. Martiel, T. Cambier, L. Blanchoin, R. Boujemaa-Paterski, and M. Théry. 2010. Nucleation geometry governs ordered actin networks structures. Nat. Mater. 9:827–832. doi:10.1038/nmat2855.

Roca-Cusachs, P., A. Del Rio, E. Puklin-Faucher, N.C. Gauthier, N. Biais, and M.P. Sheetz. 2013. Integrin-dependent force transmission to the extracellular matrix by α-actinin triggers adhesion maturation. Proc. Natl. Acad. Sci. U. S. A. 110:E1361–70. doi:10.1073/pnas.1220723110.

Sardet, C. 2015. Plankton, wonders of the drifting world. The Univeristy of Chicago Press, Chicago. 224 pp.

Schiffhauer, E.S., T. Luo, K. Mohan, E.R. Griffis, P.A. Iglesias, D.N.R. Correspondence, V. Srivastava, X. Qian, and D.N. Robinson. 2016. Mechanoaccumulative Elements of the Mammalian Actin Cytoskeleton. Curr. Biol. 26:1473–1479. doi:10.1016/j.cub.2016.04.007.

Schiller, H.B., M.-R. Hermann, J. Polleux, T. Vignaud, S. Zanivan, C.C. Friedel, Z. Sun, A. Raducanu, K.-E. Gottschalk, M. Théry, M. Mann, and R. Fässler. 2013. β1-and αv-class integrins cooperate to regulate myosin II during rigidity sensing of fibronectin-based microenvironments. Nat. Cell Biol. 15:625–636. doi:10.1038/ncb2747.

Schuppler, M., F.C. Keber, M. Kröger, and A.R. Bausch. 2016. Boundaries steer the contraction of active gels. Nat. Commun. 7:13120. doi:10.1038/ncomms13120.

Sen, S., M. Dong, and S. Kumar. 2009. Isoform-specific contributions of alpha-actinin to glioma cell mechanobiology. PLoS One. 4:e8427. doi:10.1371/journal.pone.0008427.

Shemesh, T., A.B. Verkhovsky, T.M. Svitkina, A.D. Bershadsky, and M.M. Kozlov. 2009. Role of focal adhesions and mechanical stresses in the formation and progression of the lamellipodium-lamellum interface [corrected]. Biophys. J. 97:1254–64. doi:10.1016/j.bpj.2009.05.065.

Sjöblom, B., A. Salmazo, and K. Djinović-Carugo. 2008. Alpha-actinin structure and regulation. Cell. Mol. Life Sci. 65:2688–701. doi:10.1007/s00018-008-8080-8.

Steger, G. 1998. An unbiased detector of curvilinear structures. IEEE Trans. Pattern Anal. Mach. Intell. 20:113–125. doi:10.1109/34.659930.

Sun, Z., H.-Y. Tseng, S. Tan, F. Senger, L. Kurzawa, D. Dedden, N. Mizuno, A.A. Wasik, M. Théry, A.R. Dunn, and R. Fässler. 2016. Kank2 activates talin, reduces force transduction across integrins and induces central adhesion formation. Nat. Cell Biol. 18:941–953. doi:10.1038/ncb3402.

Théry, M. 2010. Micropatterning as a tool to decipher cell morphogenesis and functions. J. Cell Sci. 123:4201–4213. doi:10.1242/jcs.075150.

Théry, M., V. Racine, M. Piel, A. Pépin, A. Dimitrov, Y. Chen, J.-B. Sibarita, and M. Bornens. 2006. Anisotropy of cell adhesive microenvironment governs cell internal organization and orientation of polarity. Proc. Natl. Acad. Sci. U. S. A. 103:19771–6. doi:10.1073/pnas.0609267103.

Thielicke, W., and E.J. Stamhuis. 2014. PIVlab – Towards User-friendly, Affordable and Accurate Digital Particle Image Velocimetry in MATLAB. J. Open Res. Softw. 2. doi:10.5334/jors.bl.

Tojkander, S., G. Gateva, A. Husain, R. Krishnan, and P. Lappalainen. 2015. Generation of contractile actomyosin bundles depends on mechanosensitive actin filament assembly and disassembly. Elife. 4:e06126. doi:10.7554/eLife.06126.

Tojkander, S., G. Gateva, and P. Lappalainen. 2012. Actin stress fibers x- assembly, dynamics and biological roles. J. Cell Sci. doi:10.1242/jcs.098087.

Tseng, Q., E. Duchemin-Pelletier, A. Deshiere, M. Balland, H. Guillou, O. Filhol, and M. Théry. 2012. Spatial organization of the extracellular matrix regulates cell-cell junction positioning. Proc. Natl. Acad. Sci. U. S. A. 109:1506–11. doi:10.1073/pnas.1106377109.

Tseng, Q., I. Wang, E. Duchemin-Pelletier, A. Azioune, N. Carpi, J. Gao, O. Filhol, M. Piel, M. Théry, and M. Balland. 2011. A new micropatterning method of soft substrates reveals that different tumorigenic signals can promote or reduce cell contraction levels. Lab Chip. 11:2231–40. doi:10.1039/c0lc00641f.

Verkhovsky, A.B., T.M. Svitkina, and G.G. Borisy. 1999. Self-polarization and directional motility of cytoplasm. Curr. Biol. 9:11–20.

Vignaud, T., L. Blanchoin, and M. Théry. 2012a. Directed cytoskeleton self-organization. Trends Cell Biol. 22:671–682. doi:10.1016/j.tcb.2012.08.012.

Vignaud, T., H. Ennomani, and M. Théry. 2014. Polyacrylamide hydrogel micropatterning. Methods Cell Biol. 120:93–116. doi:10.1016/B978-0-12-417136-7.00006-9.

Vignaud, T., R. Galland, Q. Tseng, L. Blanchoin, J. Colombelli, and M. Théry. 2012b. Reprogramming cell shape with laser nano-patterning. J. Cell Sci. 125:2134–40. doi:10.1242/jcs.104901.

Vogel, S.K., Z. Petrasek, F. Heinemann, and P. Schwille. 2013. Myosin motors fragment and compact membrane-bound actin filaments. Elife. 2:1–18. doi:10.7554/eLife.00116.

Ward, S.M.V., A. Weins, M.R. Pollak, and D.A. Weitz. 2008. Dynamic Viscoelasticity of Actin Cross-Linked with Wild-Type and Disease-Causing Mutant α-Actinin-4. Biophys. J. 95:4915–4923. doi:10.1529/BIOPHYSJ.108.131722.

Young, J.L., A.W. Holle, and J.P. Spatz. 2016. Nanoscale and mechanical properties of the physiological cell-ECM microenvironment. Exp. Cell Res. 343:3–6. doi:10.1016/j.yexcr.2015.10.037.

